# RNA helicase DDX1 regulates germinal centre selection and affinity maturation by promoting tRNA ligase activity

**DOI:** 10.1101/2025.01.10.632317

**Authors:** Rachael Kimber, Sarah Ingelsfield, Michael Screen, Fiamma Salerno, Louise Matheson, Hanneke Okkenhaug, Alex Whale, Jayalini Assalaarachchi, Melanie Stammers, Deniz Akdeniz, Simon Andrews, Claudia Ribeiro de Almeida

## Abstract

Clonal expansion of antigen-specific B-cells defines effective germinal centre responses and is key for the generation of high-affinity antibodies. While positive selection in germinal centres has been associated with anabolic metabolism and cell growth, the downstream drivers of B-cell proliferation are not well understood. Here we report that the RNA helicase DDX1 is required for germinal centre maturation and accrual of dark-zone cellularity. Upon interaction with T-follicular helper cells, DDX1-deficient B-cells upregulate c-MYC but do not clonally expand. We show that positive selection is coupled with an increase in mRNA translation, that is dependent on DDX1. DDX1 endows B-cells with the protein biosynthetic capability that is required for rapid cell proliferation. It does so by modulating the activity of the tRNA ligase complex and tRNA splicing. Our data reveal that mRNA translation efficiency is a key determinant of B-cell fitness during germinal centre responses.

## Introduction

A hallmark of adaptive immunity is the ability of B-cells to generate high affinity antibody responses following infection or immunisation. During T-cell dependent immune responses, this is achieved through formation of specialised microanatomical structures in lymphoid tissues, known as germinal centres (GCs)^1^. Once formed, GCs quickly polarize into two phenotypically and anatomically distinct compartments known as the dark zone (DZ) and the light zone (LZ). Elevated levels of the enzyme Activation Induced Cytidine Deaminase (AID) in DZ GC B-cells result in the introduction of point mutations into the variable regions of immunoglobulin heavy-chain (IgH) and light-chain (IgL) loci, a process known as Somatic Hypermutation (SHM). AID-induced mutation is targeted to sites of antigen recognition and thus enables enhancement of antibody affinity when coupled to antigen-driven selection of GC B-cells in the LZ^2, 3^. B-cells expressing newly mutated Ig on their surface as a B-cell receptor (BCR) capture antigen displayed on the surface of follicular dendritic cells (FDCs), internalise, and present the peptides on major histocompatibility complex class II (MHC II) to T-follicular helper (T_FH_) cells. Following interaction with T_FH_ cells, transient upregulation of the transcription factor c-MYC marks a small subset of positively selected B-cells, which are fated for re-entry into the DZ or differentiation into memory B-cells (MBC) and plasma cells (PC)^4, 5, 6^. DZ entrants undergo further rounds of SHM and proliferation in direct proportion to the amount of antigen they capture and present to T_FH_ cells, thereby promoting selective expansion of B-cell clones with the highest affinity for antigen^7, 8, 9^.

Clonal expansion of GC B-cells is known to be coupled to extensive changes in transcriptional and metabolic programmes, required to increase biosynthetic capacity, cell growth and proliferation. The current model proposes T_FH_-cell interactions ‘refuel’ LZ B-cells in an affinity-dependent manner making these B-cells competent to execute rapid rounds of cell division^10^. Activation of the mechanistic target of rapamycin complex 1 (mTORC1) is required early after positive selection to trigger the anabolic cell growth that sustains the proliferative burst characteristic of DZ B-cells^11^. Furthermore, mTORC1 signalling acts complementary to the transcriptional activity of c-MYC, to control cell division and metabolism^9^. Other transcription factors propagate signals received during transient T_FH_-B-cell interactions in the LZ into selective expansion of DZ B-cell clones^12, 13, 14, 15, 16^. Although protein synthesis is likely to play a key role in endowing positively selected LZ B-cells with the ability to rapidly proliferate in the DZ, the extent to which regulation of mRNA translation controls proliferation remains poorly understood^17, 18^.

Here we studied the contribution of DEAD-box RNA helicase 1 (DDX1) to humoral immune responses *in vivo*. Our previous work uncovered a requirement of DDX1 for targeting AID via a RNA G-quadruplex dependent mechanism during Class Switch Recombination (CSR), a process in which activated B-cells replace their IgH constant regions to produce alternative antibody isotypes (i.e. IgG, IgE and IgA)^19^. These findings prompted us to investigate whether DDX1 participates in AID-induced SHM in GCs. We demonstrate that DDX1-depleted B-cells fail to form functional GC structures upon immunisation. This defect was uncoupled from DDX1’s function in AID targeting and does not result from impaired SHM. Instead, we discovered a crucial role for DDX1 in maintaining B-cell clonal expansion in the DZ upon T_FH_-mediated selection. GC B-cells lacking DDX1 upregulate c-MYC but are strongly impaired in elevating the rate of protein synthesis after positive selection. Consistent with DDX1 being required for the activity of the tRNA ligase complex (tRNA-LC)^20, 21^, we observed defective maturation of intron-containing tRNAs in DDX1-deficient B-cells upon activation. Our work identifies DDX1 activity as a crucial regulator of tRNA maturation and mRNA translation during the GC response, that is essential for affinity maturation and the generation of antibodies with high antigen specificity.

## Results

### DDX1 is required during early B-cell activation and the germinal centre response

To define the role of DDX1 during humoral immune responses, we evaluated mice with conditional *Ddx1* gene deletion in naive B-cells (using *Cd23*^Cre^) or upon B-cell activation (using *Aicda*^Cre^). Mice were immunised intraperitoneally with the T-cell dependent immunogen 4-hydroxy-3-nitrophenyl (NP) coupled to keyhole limpet hemocyanin (KLH) (NP-KLH) in alum. The presence of CD38^lo^CD95^+^ GC B-cells (or alternatively CD95^+^GL7^+^PNA^+^ GC B-cells) within the spleen were characterised 7d post immunisation using flow cytometry (Fig. 1, A and B). Deletion of *Ddx1* prior to B-cell activation (*Ddx1*^fl/fl^ *Cd23*^Cre^ mice) did not affect follicular (FO, CD23^hi^CD21/35^lo^) and marginal zone (MZ, CD23^lo^CD21/35^hi^) splenic B-cell populations (Supplementary Fig. 1, A and B). However, upon immunisation, *Ddx1*^fl/fl^ *Cd23*^Cre^ mice showed a 4-fold reduction in the percentage and number of GC B-cells, compared to littermate controls (Ctrl) (Fig. 1, A and C). Corroborating our previous findings on DDX1’s contribution to CSR^19^, we detected a strongly reduced percentage of IgG1^+^ GC B-cells and retention of IgM expression in *Ddx1*^fl/fl^ *Cd23*^Cre^ GC B-cells (Fig. 1, A and D). We subsequently analysed *Ddx1*^fl/fl^ mice crossed to *Aicda*^Cre^ (referred to herein as *Ddx1*^fl/fl^ *Aicda*^Cre^), where *Ddx1* deletion occurs once AID expression is induced following B-cell activation. Notably, the percentage and number of GC B-cells present in *Ddx1*^fl/fl^ *Aicda*^Cre^ mice 7d post-immunisation were comparable to littermate controls (Ctrl) (Fig. 1, B and E). Likewise, we detected similar proportions of IgG1^+^ GC B-cells in *Ddx1*^fl/fl^ *Aicda*^Cre^ and Ctrl mice (Fig. 1, B and F). However, when analysing *Ddx1*^fl/fl^ *Aicda*^Cre^ mice 10d post-immunisation, we observed a 2-fold reduction in the number of GC B-cells (Fig. 1E). These results indicated that the timing of *Ddx1* deletion during B-cell immune responses differentially impacts CSR and the GC reaction.

**Figure 1.**
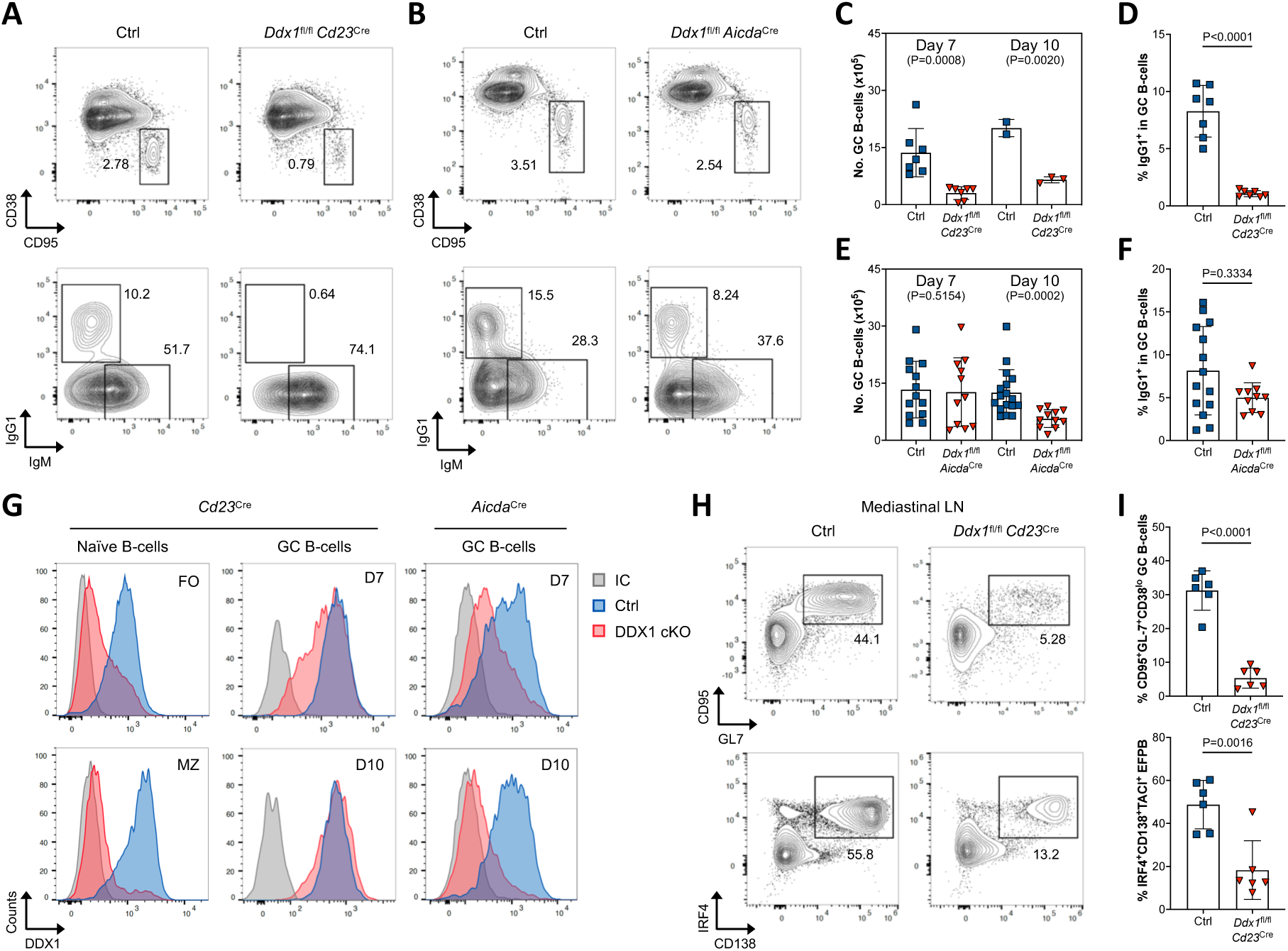
Depletion of Ddx1 at different steps during B-cell activation uncouples defects on CSR from germinal centre formation. **a,b**, Representative flow cytometry plots of GC B-cells (CD38^lo^CD95^+^) within the total CD19^+^B220^+^ B-cell population in *Ddx1*^fl/fl^ *Cd23*^Cre^ (**a**), *Ddx1*^fl/fl^ *Aicda*^Cre^ (**b**) and littermate control (Ctrl) mice, 7 days (7d) post NP-KLH immunisation. **c**, Quantification of the total number of GC B-cells in Ctrl and *Ddx1*^fl/fl^ *Cd23*^Cre^ mice 7d and 10d post-immunisation. **d**, Quantification of the frequency of IgG1^+^ GC B-cells in Ctrl and *Ddx1*^fl/fl^ *Cd23*^Cre^ mice 7d post-immunisation. **e**, Quantification of the total numbers of GC B-cells in Ctrl and *Ddx1*^fl/fl^ *Aicda*^Cre^ mice 7d and 10d post-immunisation. **f**, Quantification of the frequency of IgG1^+^ GC B-cells in Ctrl and *Ddx1*^fl/fl^ *Aicda*^Cre^ mice 7d post-immunisation. GC B-cells quantified in (**c**) and (**e**) were defined as CD95^+^GL-7^+^PNA^+^ or CD38^lo^CD95^+^. Each datapoint represents an individual mouse, bar represent mean ± SD. Data is from 3-4 independent experiments. The *P* values were generated by performing an unpaired, two-sided Student’s t-test on logarithmically transformed data. **g**, Representative histograms for intracellular DDX1 protein levels measured by flow cytometry in FO and MZ B-cells from non-immunised mice, and GC B-cells 7d and 10d post-immunisation. Signal obtained using an anti-DDX1 monoclonal antibody is compared to response for isotype control (IC). **h**, Representative flow cytometry plots of GC B-cells (CD95^+^GL-7^+^) in CD138^-^IgD^-^CD19^+^ B-cells (top) and EFPB (IRF4^+^CD138^+^) in IgD^-^CD19^+^ B-cells (bottom) in mediastinal lymph nodes (mLN) from Ctrl and *Ddx1*^fl/fl^ *Cd23*^Cre^ mice, 8d post HKx31 Influenza A infection. **i**, Quantification of the frequency of GC B-cells (top) and EFPB (bottom) defined as CD95^+^GL-7^+^CD38^lo^ and IRF4^+^CD138^+^TACI^+^, respectively. Each datapoint represents an individual mouse, bar represent mean ± SD. Data is from 2 independent experiments. The *P* values were generated by performing an unpaired, two-sided Student’s t-test on logarithmically transformed data.

We evaluated the kinetics of DDX1 protein depletion in both models using intracellular flow cytometry staining. In *Ddx1*^fl/fl^ *Cd23*^Cre^ mice, DDX1 protein levels were strongly reduced in naïve FO and MZ B-cells (Fig. 1G). Remarkably, GC B-cells present in *Ddx1*^fl/fl^ *Cd23*^Cre^ mice retained levels of DDX1 protein comparable to Ctrl mice, indicating strong selection for cells that escaped *Ddx1* gene deletion. These results suggest that expression of DDX1 is required for establishing GC B-cells (Fig. 1G and Supplementary Fig. 1D). In *Ddx1*^fl/fl^ *Aicda*^Cre^ mice, we observed partial depletion of DDX1 protein 7d post-immunisation (Fig. 1G). However, DDX1 protein levels are further reduced as the GC response matures at d10 (Fig. 1G and Supplementary Fig. 1E). In summary, our results indicate that in *Ddx1*^fl/fl^ *Aicda*^Cre^ mice sufficient DDX1 protein expression is retained at earlier timepoints of the immune response enabling CSR and GC seeding. As the immune response progresses, efficient DDX1 depletion results in a loss of GC B-cells in *Ddx1*^fl/fl^ *Aicda*^Cre^ mice, possibly due to defects in GC maturation and/or maintenance.

To determine the effect of DDX1 depletion in the response of B-cells to polyclonal antigen exposure, we infected *Ddx1*^fl/fl^ *Cd23*^Cre^ mice intranasally with HKx31 Influenza A. Although, *Ddx1*^fl/fl^ *Cd23*^Cre^ and littermate controls showed comparable disease progression as measured by weight loss (Supplementary Fig. 1, F and G), a strong reduction in CD95^+^GL7^+^ GC B-cells was observed in the mediastinal lymph node (mLN) and spleen of *Ddx1*^fl/fl^ *Cd23*^Cre^ mice 8d post-infection (Fig. 1, H and I and Supplementary Fig. 1, H and I). Influenza A infection also drives a large extrafollicular plasmablast (EFPB) response, particularly within the mLN, enabling the evaluation of early activated B-cell populations^22^. Strikingly, only a small percentage of IRF4^+^CD138^+^ EFPB were observed in *Ddx1*^fl/fl^ *Cd23*^Cre^ mice, highlighting that DDX1 depletion restricts initial activated B-cell responses (Fig. 1, H and I and Supplementary Fig. 1, H and I). We observed incomplete DDX1 protein depletion in EFPB and GC B-cell populations from *Ddx1*^fl/fl^ *Cd23*^Cre^ mice upon influenza infection, with a strong selection for EFPB retaining substantial levels of DDX1 protein in the mLN (Supplementary Fig. 1J). Overall, this indicates that DDX1 expression is key to early B-cell activation, the generation of EFPB, and to maintain effective GC B-cell responses.

### Loss of dark zone in germinal centres of *Ddx1*^fl/fl^ *Aicda*^Cre^ mice

We evaluated if the reduction in GC B-cells observed in *Ddx1*^fl/fl^ *Aicda*^Cre^ mice was due to impaired differentiation of DZ (CXCR4^hi^CD86^lo^) and LZ (CXCR4^lo^CD86^hi^) subsets at d10 post-immunisation using flow cytometry (Fig. 2A). Strikingly, we observed a significant reduction in the percentage of DZ B-cells in *Ddx1*^fl/fl^ *Aicda*^Cre^ mice, and a concomitant skewing towards a higher proportion of LZ B-cells, resulting in reduced DZ/LZ ratios (Fig. 2, B-D). Importantly, we did not observe defects in the proportion of DZ and LZ B-cells in *Ddx1*^fl/wt^ *Aicda*^Cre^ or *Ddx1*^wt/wt^ *Aicda*^Cre^ mice, suggesting that *Ddx1* gene dosage or the presence of the *Aicda*^Cre^ transgene do not impact DZ and LZ B-cell differentiation (Fig. 2A-D and Supplementary Fig. 2A and B).

**Figure 2.**
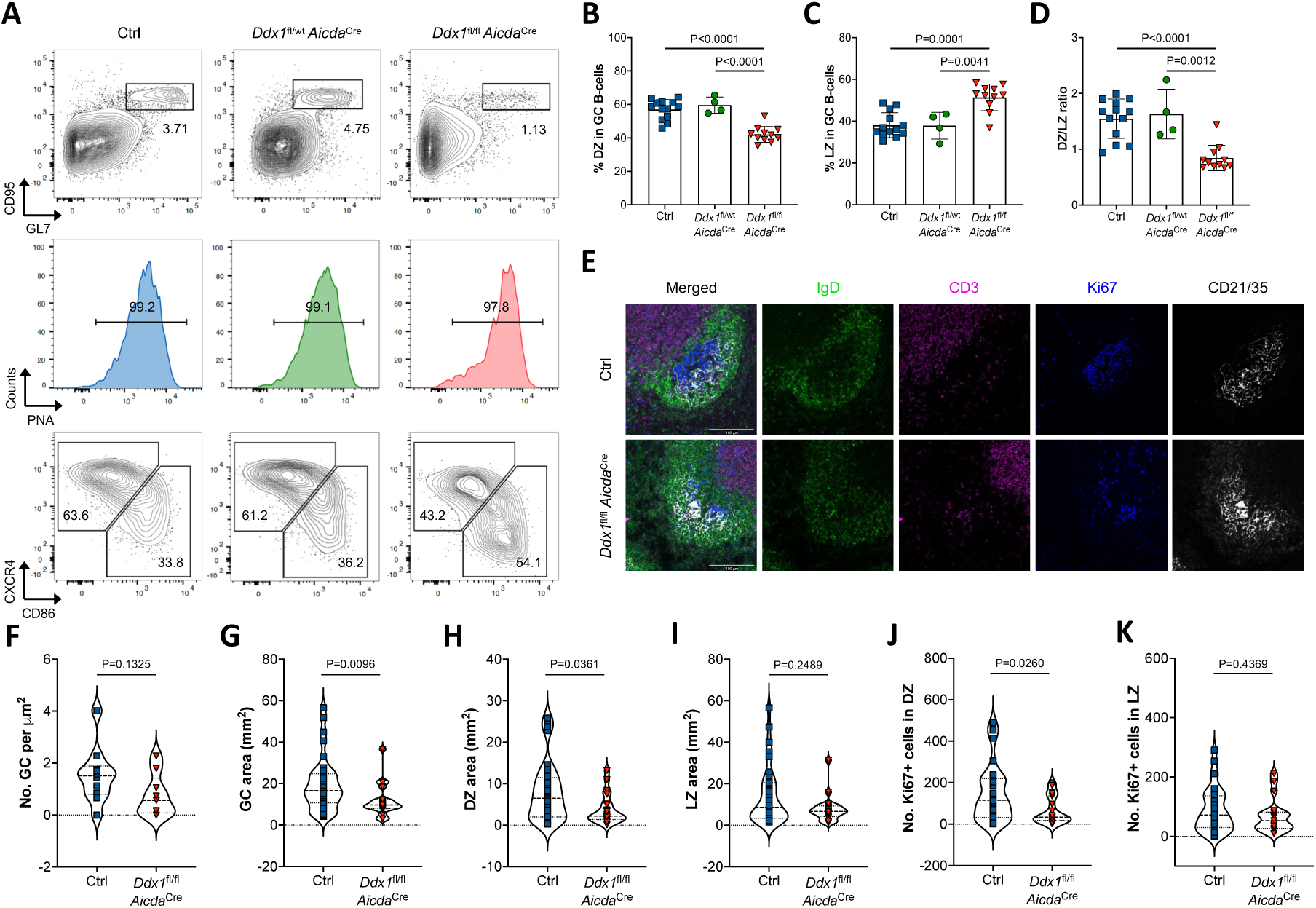
DDX1-depleted Germinal Centres are reduced in size due to a collapse of the dark zone. **a,** Representative flow cytometry plots of GC B-cells (CD95^+^GL-7^+^PNA^+^) within the total CD19^+^B220^+^ B-cell population in spleens from Ctrl, *Ddx1*^fl/wt^ *Aicda*^Cre^ and *Ddx1*^fl/fl^ *Aicda*^Cre^ mice, 10d post NP-KLH immunisation. DZ and LZ B-cells were defined based on the reciprocal expression of the surface markers CXCR4 and CD86. **b-d**, Quantification of the frequency of CXCR4^hi^CD86^lo^ DZ (**b**) and CXCR4^lo^CD86^hi^ (**c**) LZ B-cells, and DZ/LZ ratios (**d**). Each datapoint represents an individual mouse, bar represent mean ± SD. Data is from 4 independent experiments. The *P* values were generated by performing a one-way ANOVA on logarithmically transformed data, with Holm-Sidak correction for multiple testing. **e,** Representative confocal images of GC structures in splenic sections from Ctrl and *Ddx1*^fl/fl^ *Aicda*^Cre^ mice, 10d post immunisation. GC structures were identified as adjacent clusters of Ki67^+^ (blue) B-cells and CD21/35^+^ (white) follicular dendritic cells within IgD^+^ B-cell follicles (green). T-cell zone are marked as CD3^+^ (purple). The scale bar represents 100 μm. **f**, Quantification of the number of GC per nm^2^ of splenic section. **g-i**, Quantification of GC area (**g**) and DZ (**h**) and LZ (**i**) areas within individual GC structures. **j-k**, Quantification of the number of Ki67^+^ cells present in DZ (**j**) and LZ (**k**) areas. Dashed lines represent the median and quartiles. The *P* values were generated by performing an unpaired, two-sided Mann-Whitney *U* test.

We determined whether the preferential loss of DZ B-cells was due to biased depletion of DDX1 mediated by the *Aicda*^Cre^ model, as AID expression is known to be greater in DZ B-cells actively undergoing SHM^17^. We detected comparable DDX1 protein expression within DZ and LZ B-cells in Ctrl mice, with similar DDX1 depletion observed in both subsets (Supplementary Fig. 2C). We also determined the frequency of class-switched GC B-cells at d10 post-immunisation. Although the frequency of class-switched B-cells in GC from Ctrl and *Ddx1*^fl/fl^ *Aicda*^Cre^ mice was comparable at d7 (Fig. 1F), we observed a decline in IgG1^+^ B-cells and retention of IgM^+^ B-cells in *Ddx1*^fl/fl^ *Aicda*^Cre^ mice as the GC matured (Supplementary Fig. 2, D and E). We confirmed the reduction in DDX1 protein levels was comparable in IgG1^+^ and IgM^+^ GC B-cells, demonstrating the presence of class-switched B-cells in the GC is not a consequence of incomplete DDX1 protein depletion (Supplementary Fig. 2F). Our results indicate that in *Ddx1*^fl/fl^ *Aicda*^Cre^ mice GC B-cells are impaired in the acquisition of the DZ phenotype and skewed towards IgM over IgG1 isotypes.

To analyse the number and morphology of GC structures we performed immunofluorescence (IF) staining and confocal imaging in splenic sections from *Ddx1*^fl/fl^ *Aicda*^Cre^ and Ctrl mice, 10d post-immunisation. Clusters of Ki67^+^ proliferating B-cells adjacent to CD21/35^+^ FDCs indicated the presence of GCs, and these were typically surrounded by naive B-cells within splenic follicles (IgD^+^) and located close to the T-cell zone (CD3^+^) (Fig. 2E). Prior to IF analyses, we validated the use of Ki67 marker by observing comparable intracellular staining intensities between *Ddx1*^fl/fl^ *Aicda*^Cre^ and Ctrl mice using flow cytometry (Supplementary Fig. 2G). To enable quantification of the number and size of GC areas, alongside the number of Ki67^+^ B-cells, we developed an automated analysis pipeline (Supplementary Fig. 2, H and I; see Methods). We observed a similar number of GC structures per μm^2^ of splenic section between *Ddx1*^fl/fl^ *Aicda*^Cre^ and Ctrl mice (Fig. 2F). In contrast, we observed a significant reduction in size of the GC area in *Ddx1*^fl/fl^ *Aicda*^Cre^ mice (Fig. 2G). Further characterisation of GC structures as DZ and LZ areas, indicated *Ddx1*^fl/fl^ *Aicda*^Cre^ mice have GCs with smaller DZs and a reduced number of Ki67^+^ GC B-cells, specifically in the DZ (Fig. 2, H-K). These results support our earlier observation that in *Ddx1*^fl/fl^ *Aicda*^Cre^ mice GC formation is unaffected (Fig. 1G). However, these GC structures appeared to be halted in their maturation, with fewer Ki67^+^proliferating B-cells leading to a specific loss of the DZ compartment.

### DDX1 is required for affinity maturation of the antibody response

To determine the extent to which the collapse of GC maturation and loss of the DZ impacted the function of the GC, we assessed the abundance of NP-specific antibodies in sera from *Ddx1*^fl/fl^ *Aicda*^Cre^ and Ctrl mice following NP-KLH immunisation. Quantification of NP-binding antibodies was performed by ELISA utilising NP_9_ for IgM quantification, or broad-affinity NP_27_ and high-affinity NP_2_ antigens for IgG1 quantification. We observed no differences in NP-binding IgM antibodies between *Ddx1*^fl/fl^ *Aicda*^Cre^ and Ctrl mice (Fig. 3A and Supplementary Fig. 3A). In contrast, a significant reduction in titres for broad-affinity and high-affinity IgG1 antibodies was observed in *Ddx1*^fl/fl^ *Aicda*^Cre^ mice (Fig. 3B and C; Supplementary Fig. 3B and C). In accordance with efficient affinity maturation, an increase in the ratio of high-affinity (NP_2_ binding) compared to broad-affinity (NP_27_ binding) IgG1 antibodies was observed over time in Ctrl mice (i.e., increased NP_2_/NP_27_ ratios; Fig. 3D). Notably, *Ddx1*^fl/fl^ *Aicda*^Cre^ mice showed impaired NP-specific affinity maturation with NP_2_/NP_27_ ratios low at all time-points (Fig. 3D). These results indicate that DDX1 is required for the generation of antigen-specific antibodies and their affinity maturation.

**Figure 3.**
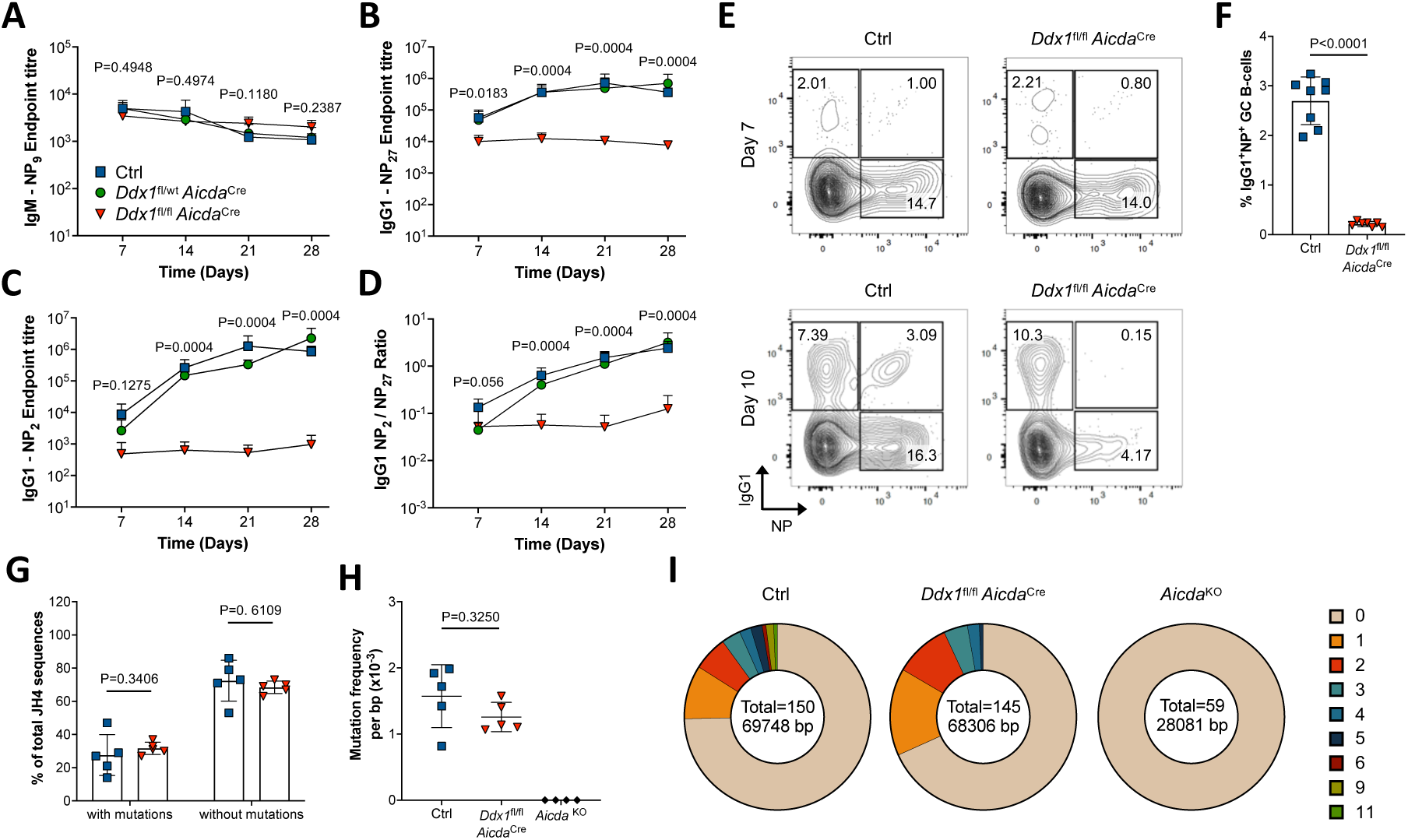
*Ddx1*^fl/fl^ *Aicda*^Cre^ mice are unable to generate high-affinity antibody responses. **a-c**, Quantification of endpoint titres for IgM-NP_9_ antibodies (**a**), broad-affinity IgG1-NP_27_ antibodies (**b**) and high-affinity IgG1-NP_2_ antibodies (**c**) in the serum of Ctrl, *Ddx1*^fl/wt^ *Aicda*^Cre^ and *Ddx1*^fl/fl^ *Aicda*^Cre^ mice at multiple time-points post-immunisation with NP-KLH. **d**, Quantification of the ratio of IgG1-NP_2_ over IgG1-NP_27_ antibody. Each datapoints represents the mean ± S.D. The *P* values were generated by performing a one-way ANOVA on logarithmically transformed data, with Holm-Sidak correction for multiple testing. **e**, Representative flow cytometry plots of IgG1 expression and NP_29_-PE binding in splenic GC B-cells from Ctrl and *Ddx1*^fl/fl^ *Aicda*^Cre^ mice 7d (top) and 10d (bottom) post-immunisation. **f**, Quantification of the frequency of IgG1^+^NP^+^ GC B-cells 10d post-immunisation, gated in (**e**). **g**, Quantification of the frequency of J_H_4 intron sequences assessed containing 1 or more mutations. **h**, Quantification of mutational frequency per basepair (bp) in J_H_4 intron sequences. Each datapoint represents an individual mouse, bar represent mean ± S.D. Data is from 2 independent experiments. The *P* values were generated by performing an unpaired, two-sided Student’s t-test on logarithmically transformed data. **i**, Fraction of clones with indicated number of mutations represented as section of a circle. Total number of clones assessed and total number of basepairs quantified are indicated.

To ascertain if the reduction in antigen-specific antibodies stems from an inability of *Ddx1*^fl/fl^ *Aicda*^Cre^ mice to generate NP-specific B-cells during GC responses, NP_29_-PE binding to IgG1^+^ GC B-cells was measured by flow cytometry (Fig. 3E). We readily detected an increase in the percentage of IgG1^+^NP^+^ GC B-cells in Ctrl mice, as the GC response to NP-KLH matures from d7 to d10 (Fig. 3E). Most notably, while these cells could be detected in both *Ddx1*^fl/fl^ *Aicda*^Cre^ and Ctrl mice at d7, DDX1-depletion resulted in an almost complete abolishment of IgG1^+^NP^+^ GC B-cells at d10 (Fig. 3, E and F). This demonstrates that *Ddx1*^fl/fl^ *Aicda*^Cre^ mice have equivalent initial responses to NP-KLH immunisation as Ctrl mice but as the GC reaction matures, DDX1 depletion severely impairs the expansion of IgG1^+^ NP-specific B-cells.

To further characterise the impact of DDX1-depletion on GC output we examined the frequency and phenotype of MBC. MBC subsets were defined based on the expression of CD38 and PD-L2, while co-expression of CD80 and CD73 was used to identify a sub-population of MBCs with highly mutated BCRs arising from the GC^24, 25^. We observed comparable percentages and total numbers of CD38^int^PD-L2^+^ MBC between *Ddx1*^fl/fl^ *Aicda*^Cre^ and Ctrl mice (Supplementary Fig. 3, D and E). Further stratification by IgG1 and IgM expression highlighted a significant reduction in percentage of IgG1^+^ MBC in *Ddx1*^fl/fl^ *Aicda*^Cre^ mice, while the percentage of IgM^+^ MBC was similar to Ctrl mice (Supplementary Fig. 3D, F and G). CD80 and CD73 expression demonstrated a bias for *Ddx1*^fl/fl^ *Aicda*^Cre^ mice to have a higher percentage of MBC that were negative for both markers (Supplementary Fig. 3, D and H). Inversely, the percentage of CD80^+^CD73^+^ MBC in *Ddx1*^fl/fl^ *Aicda*^Cre^ mice was reduced compared to Ctrl mice (Supplementary Fig. 3, D and H). This indicates that GC-specific depletion of DDX1 does not impact on the quantity of MBC generated. Rather, the ability to generate class-switched MBC and especially those with a phenotype associated with high-affinity BCRs required DDX1.

Taken together, these analyses demonstrate that DDX1 depletion in the GC does not restrict the output of antigen-specific IgM antibodies or IgM^+^ MBC. However, affinity maturation of class-switched GC B-cells and the antibody response, as well as the generation of MBC populations with increased affinity towards antigen, require DDX1 expression in B cells.

### DDX1 deficiency impacts on germinal centre maturation independently of AID activity

We determined whether the defects in high-affinity, antigen-specific immune responses in *Ddx1*^fl/fl^ *Aicda*^Cre^ mice was due to an impairment in SHM. To address this, we sorted IgG1^+^ GC B-cells 10d post-immunisation and analysed the sequence of J_H_4 intron amplicons, a variable region surrogate that enables quantification of the mutational rate of AID independently of positive selection^26^. Comparable numbers of sequences containing mutations were recovered from Ctrl and *Ddx1*^fl/fl^ *Aicda*^Cre^ mice (48 out of 148, and 42 out of 152, respectively) (Fig. 3H). Although not significant, *Ddx1*^fl/fl^ *Aicda*^Cre^ mice demonstrated a reduced mutational frequency compared to Ctrl, which was on average 1.77×10^4^/bp and 1.22×10^4^/bp, respectively, whereas only germline sequences were recovered from *Aicda*^KO^ mice as expected (Fig. 3I). Comparison of the number of mutations per sequence highlighted that *Ddx1*^fl/fl^ *Aicda*^Cre^ mice had a higher proportion of sequences with only one or two mutations, whereas Ctrl mice were able to produce sequences with up to eleven mutations (Fig. 3J). This indicates that although GC B-cells in *Ddx1*^fl/fl^ *Aicda*^Cre^ mice can undergo SHM, they are unable to generate clones with a high mutational load, resulting in a trend for a reduced overall mutational rate.

To formally establish that DDX1-mediated loss of DZ GC B-cells is independent of its known role in AID activity^19^, we crossed *Ddx1*^fl/fl^ *Aicda*^Cre^ mice on to an *Aicda*^KO^ background. Upon immunisation *Aicda*^KO^ mice show enlarged GC and LZ compartments due to lower rates of apoptosis and defects in MHCII downregulation when transitioning to the DZ^27, 28, 29^. We observed GC hyperplasia with a significant increase in percentage and total numbers of GC B-cells in *Aicda*^KO^ mice (Supplementary Fig. 4, A and B). Notably, the response for *Ddx1*^fl/fl^ *Aicda*^Cre^ *Aicda*^KO^ mice was significantly reduced compared to that of *Aicda*^KO^ Ctrl mice, with a 2.5-fold decrease in the number of GC B-cells detected (Supplementary Fig. 4B). We observed a significant decrease in the percentage of DZ GC B-cells in *Aicda*^KO^ mice (Supplementary Fig. 4, C and D). Strikingly, depletion of DDX1 in *Aicda*^KO^ mice resulted in further reduction in the DZ compartment compared to DDX1-sufficient counterparts, with an additional increase in the percentage of LZ GC B-cells (Supplementary Fig. 4, C-F). This demonstrates that DDX1 function in GC maturation and the generation of high-affinity, antigen-specific immune responses are independent of AID activity.

### DDX1 is dispensable for c-MYC expression by B cells upon T-cell mediated selection

A key aspect of the GC response is the ability of GC B-cells, upon acquisition of antigen, to interact with T_FH_ cells and undergo T-cell mediated selection in the LZ. Positive selection signals determine the extent of cell division and hypermutation GC B-cells undergo in the DZ^8^, with the amount of c-MYC protein expression in LZ B-cells directly correlating with the level of B-cell expansion^4, 5, 9^. We addressed whether defects in T_FH_-mediated selection and/or c-MYC upregulation in *Ddx1*^fl/fl^ *Aicda*^Cre^ mice were behind the collapse of the DZ, by crossing *Ddx1*^fl/fl^ *Aicda*^Cre^ mice with *Myc*^GFP/GFP^ mice^30^. MYC-eGFP^+^ cells were detected in the GC B-cell compartment, with an increase in the percentage of positively selected cells observed in *Ddx1*^fl/fl^ *Aicda*^Cre^ *Myc*^GFP/GFP^ mice compared to Ctrl *Myc*^GFP/GFP^ mice (Fig. 4, A and B). Within the MYC-eGFP^+^ GC B-cell population we observed similar proportions of LZ and DZ B-cells in Ctrl *Myc*^GFP/GFP^ mice, likely due to increased sensitivity of spectral flow cytometry enabling the detection of low MYC-eGFP levels in DZ B-cells (Fig. 4C). However, MYC-eGFP^+^ GC B-cells in *Ddx1*^fl/fl^ *Aicda*^Cre^ *Myc*^GFP/GFP^ mice predominantly retain a LZ phenotype, with fewer cells in the DZ compartment (Fig. 4C). To account for differences in the level of MYC-eGFP signal between genotypes, we performed similar analysis in MYC-eGFP^low^ and MYC-eGFP^high^ GC B-cells and confirmed an increased presence of LZ MYC-eGFP^+^ cells in *Ddx1*^fl/fl^ *Aicda*^Cre^ *Myc*^GFP/GFP^ mice (Supplementary Fig. 4G). Nevertheless, the intensity of MYC-eGFP signal was equally decreased upon the transition of GC B-cells from LZ to the DZ when comparing Ctrl *Myc*^GFP/GFP^ and *Ddx1*^fl/fl^ *Aicda*^Cre^ *Myc*^GFP/GFP^ mice (Fig. 4D). Thus, DDX1 expression in GC B-cells is dispensable for c-MYC induction following T-cell mediated selection.

**Figure 4.**
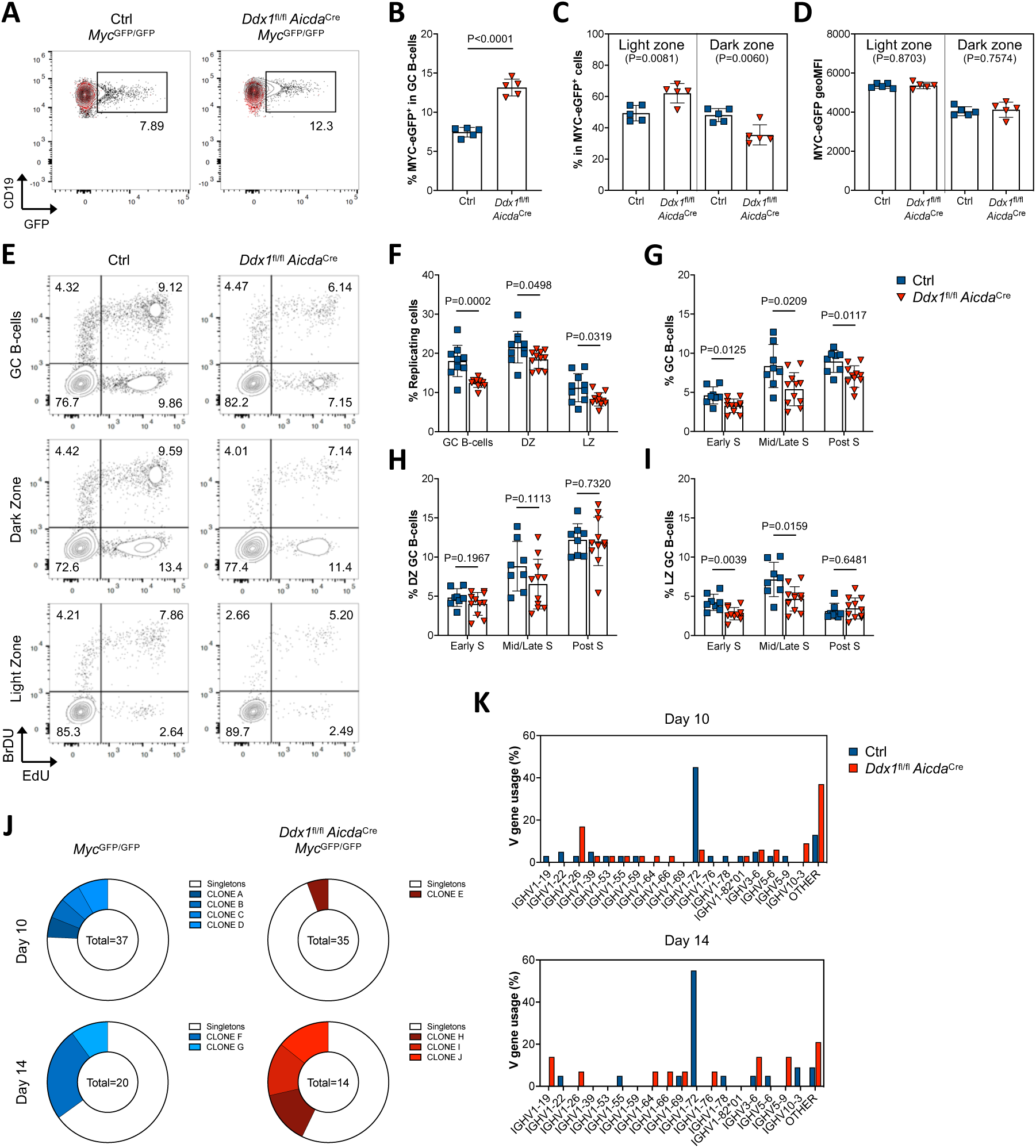
Defects in B-cell proliferation hinder clonal expansion in DDX1-deficient germinal centres. **a**, Representative flow cytometry plots of MYC-eGFP expression in GC B-cells from Ctrl *Myc*^GFP/GFP^ and *Ddx1*^fl/fl^ *Aicda*^Cre^ *Myc*^GFP/GFP^ mice, 10d post NP-KLH immunisation. An overlay of MYC-eGFP negative GC B-cells is shown in red. **b**, Quantification of the frequency of MYC-eGFP^+^ GC B-cells gated in (**a**). **c**, Quantification of the frequency of DZ and LZ subsets within MYC-eGFP^+^ GC B-cell population. **d**, Quantification of geoMFI for the MYC-eGFP reporter protein in MYC-eGFP^+^ LZ and DZ B-cells. Each datapoint represents an individual mouse, bar represent mean ± SD. Data is from one experiment. The *P* values were generated by performing an unpaired, two-sided Student’s t-test on logarithmically transformed data. **e**, Representative flow cytometry plots of EdU and BrdU incorporation in total GC, DZ and LZ B-cells in spleens from Ctrl and *Ddx1*^fl/fl^ *Aicda*^Cre^ mice, 10d post immunisation. **f**, Quantification of the frequency of replicating (EdU^+^) cells. **g-i**, Quantification of the frequency of cells in early S-phase (EdU^-^BrdU^+^), mid/late S-phase (EdU^+^BrdU^+^) and post S-phase (EdU^+^BrdU^-^). Each datapoint represents an individual mouse, bar represent mean ± S.D. Data is from 3 independent experiments. The *P* values were generated by performing an unpaired, two-sided Student’s t-test on logarithmically transformed data. **j**, Pie charts showing the distribution of unique (singletons) and expanded VDJ-Ighg1 sequences obtained from splenic MYC-eGFP^+^IgG1^+^ GC B cells sorted from Ctrl *Myc*^GFP/GFP^ and *Ddx1*^fl/fl^ *Aicda*^Cre^ *Myc*^GFP/GFP^ mice, 10d and 14d post immunisation. Data represents 1-2 mice per group and each segment represents a unique clone, with the total number of analysed sequences indicated in the centre of each chart. Note that only 2 unique sequences were recovered for each expanded clone from *Ddx1*^fl/fl^ *Aicda*^Cre^ *Myc*^GFP/GFP^ mice at 14d. **k**, Quantification of the frequency of V_H_ region usage in VDJ-Ighg1 sequences. Clones using different D_H_ and J_H_ regions were pooled and only V_H_ genes present in more than one dataset are depicted; ‘other’ denotes V_H_ genes only present in one dataset.

Programmed cellular death by apoptosis is at the core of GC clonal selection mechanisms and an important determinant of affinity maturation^28, 31^. Positive selection signals in the LZ provide protection from apoptosis whereas, unproductive mutations during SHM, such as the introduction of stop codons, results in increased apoptosis in the DZ^31, 32^. We determined whether GC defects observed in *Ddx1*^fl/fl^ *Aicda*^Cre^ mice are driven by apoptosis by measuring the levels of activated caspase-3 using flow cytometry. As expected, minimal activated caspase-3 signal was detected in non-GC B-cells whereas, an enhanced signal was observed in GC B-cells (Supplementary Fig. 4H). We observed no differences in the percentage of activated caspase-3 within total GC, DZ or LZ B-cells in *Ddx1*^fl/fl^ *Aicda*^Cre^ mice compared to Ctrl mice (Supplementary Fig. 4I). This indicates that loss of high-affinity, antigen-specific GC B-cells in DDX1 deficient mice is not due to increased apoptosis.

### DDX1 is required for clonal expansion of germinal centre B-cells

We determined whether *Ddx1*^fl/fl^ *Aicda*^Cre^ GC B-cells were able to undergo cell division and convert T_FH_-mediated positive selection cues into the proliferative burst required to populate the DZ. We used dual pulse labelling with the thymidine analogues 5-Ethynyl-2’-deoxyuridine (EdU) and bromodeoxyuridine (BrdU) to evaluate cell-cycle progression (Fig. 4E). We used EdU incorporation as a measure of GC B-cell replication and observed that while a greater proportion of EdU^+^ GC B-cells are within the DZ compartment in Ctrl mice, EdU incorporation was significantly decreased in both DZ and LZ subsets in *Ddx1*^fl/fl^ *Aicda*^Cre^ mice (Fig. 4F). This indicates that DDX1 is required for DZ and LZ B-cells to undergo DNA replication during S-phase. When further stratified, actively cycling GC B-cells can be classed into early, mid/late and post-S-phases based on BrdU incorporation in combination with EdU (Fig. 4E). We observed a significant decrease in GC B-cells in all S-phases in *Ddx1*^fl/fl^ *Aicda*^Cre^ mice compared to Ctrl mice (Fig. 4G). Dual labelling with EdU and BrdU also identified DZ B-cells as mostly present in the post S-phase of the cell-cycle, having finished DNA synthesis during the BrdU pulse period due to their high proliferative nature (Fig. 4H). This contrasts with LZ B-cells where most replicating cells were in mid/late S-phase (Fig. 4I). Although there was a significant reduction in EdU incorporation in DZ B-cells, we did not observe significant differences in S-phase progression in DZ B-cells from *Ddx1*^fl/fl^ *Aicda*^Cre^ mice compared to Ctrl mice (Fig. 4H). In contrast, there was a significant reduction of LZ B-cells in early and mid/late S-phase in *Ddx1*^fl/fl^ *Aicda*^Cre^ mice (Fig. 4I). Taken together, our results indicate that DDX1-deficient GC B-cells are impaired in S-phase entry and progression already in the LZ. These cells seem to restore their ability to complete DNA replication when transition to the DZ occurs.

We next sought to determine whether defects in GC B-cell proliferation hinder expansion of antigen-specific B-cell clones. We sequenced VDJ-Ighg1 amplicons as described by Von Boehmer et al., 2016^33^ from single MYC-eGFP^+^IgG1^+^ GC B cells sorted from Ctrl *Myc*^GFP/GFP^ and *Ddx1*^fl/fl^ *Aicda*^Cre^ *Myc*^GFP/GFP^ mice 10d and 14d post NP-KLH immunisation. Our analysis demonstrated that the magnitude of clonal diversity was impaired in GCs lacking DDX1; 10d post-immunisation, most clones recovered from *Ddx1*^fl/fl^ *Aicda*^Cre^ *Myc*^GFP/GFP^ mice have a unique VDJ sequence (singletons), while almost 25% of clones recovered in Ctrl *Myc*^GFP/GFP^ mice were expanded (Fig. 4J). After a further 4d the total expansion rate was more comparable with 35% and 43% of clones expanded in Ctrl *Myc*^GFP/GFP^ and *Ddx1*^fl/fl^ *Aicda*^Cre^ *Myc*^GFP/GFP^ mice, respectively (Fig. 4J). However, in Ctrl *Myc*^GFP/GFP^ mice, one clone had dominated the response and constituted 25% of the population, whereas expansion was limited to 14% in each of the three clones observed for *Ddx1*^fl/fl^ *Aicda*^Cre^ *Myc*^GFP/GFP^ mice (Fig. 4J).

To further understand if DDX1-depletion impaired the selection of antigen-specific clones, we inspected V-region usage between genotypes. In Ctrl *Myc*^GFP/GFP^ mice the most abundant V-region gene utilised was IGHV1-72 (Fig. 4K), which is the canonical sequence known to respond to NP and dominate the response in C57BL/6 mice^34^. In stark contrast, less than 6% of GC B-cells from *Ddx1*^fl/fl^ *Aicda*^Cre^ *Myc*^GFP/GFP^ mice had BCRs with the IGHV1-72 region at 10d post-immunisation and we were unable to recover any clones utilising this gene segment by 14d (Fig. 4K). Additionally, alternative clones (namely, IGHV1-53 and IGHV6-3) with NP-binding capacity, were detected in positively selected GC B-cells from Ctrl *Myc*^GFP/GFP^ mice (Fig. 4K, Supplementary Table 1 and 2); whereas, DDX1-deficient MYCeGFP^+^IgG1^+^ GC B-cells expressed Ig genes with minimal NP-binding e.g. IGHV1-69 (Fig. 4K, Supp Table 1 and 2)^35^. Almost two-thirds of the IGHV1-72 clones detected in Ctrl *Myc*^GFP/GFP^ mice show the characteristic tryptophan (W) to leucine (L) substitution at position 33 (W33L) and lysine (K) to arginine (R) substitution at position 59 (K59R) in their sequences; both mutations known to enhance affinity for NP^36^. However, the few IGHV1-72 clones obtained from *Ddx1*^fl/fl^ *Aicda*^Cre^ *Myc*^GFP/GFP^ GC B-cells were germline (Supplementary Table 1 and 2). Taken together, this demonstrates that even though GC B-cells in *Ddx1*^fl/fl^ *Aicda*^Cre^ *Myc*^GFP/GFP^ mice can undergo SHM, GC are unable to expand antigen-specific clones and increase their mutational load to improve antigen affinity.

### The transcriptional programme of c-MYC^+^ germinal center B-cells is not dependent on DDX1

To elucidate further the role of DDX1 in GC responses, we used single-cell RNA-Seq (scRNA-Seq) to profile gene expression in MYC-eGFP^+^ GC B-cells sorted from the spleens of Ctrl *Myc*^GFP/GFP^ and *Ddx1*^fl/fl^ *Aicda*^Cre^ *Myc*^GFP/GFP^ mice, 10d post-immunisation (Supplementary Fig. 5A). Following quality control filtering, a total of 204 cells were used for subsequent analyses. To validate our approach, we compared the index-sorted mean fluorescence intensity (MFI) for GFP in each individual cell across the two genotypes (Fig. 5A). Although a trend for a higher GFP MFI was observed in Ctrl *Myc*^GFP/GFP^ samples, with the detection of a few cells with very high levels of expression, we observed no significant difference in the GFP signal compared to *Ddx1*^fl/fl^ *Aicda*^Cre^ *Myc*^GFP/GFP^ samples (Fig. 5A). To understand the proportions of LZ and DZ B-cells identified, we scored individual cells based on transcripts defined by Victora et al., 2012^37^. When single cells were plotted based on their LZ and DZ scores, we observed a higher percentage of cells scored as LZ than DZ, particularly in *Ddx1*^fl/fl^ *Aicda*^Cre^ *Myc*^GFP/GFP^ mice (Fig. 5, B and C). To determine the accuracy of LZ and DZ scoring we overlayed the index expression data of phenotypic markers CXCR4, CD86 and CD83 (Supplementary Fig. 5B). Cells defined as DZ based on gene expression had a trend for higher expression of CXCR4, with lower expression observed in LZ cells (Supplementary Fig. 5B). However, CD83 and CD86 protein expression patterns were less conclusive (Supplementary Fig. 5B). This analysis is consistent with our previous findings of a reduction of DZ B-cells within the MYC-eGFP^+^ subset following DDX1-depletion, with CXCR4 expression correlating with gene expression changes characterising the two main GC populations.

**Figure 5.**
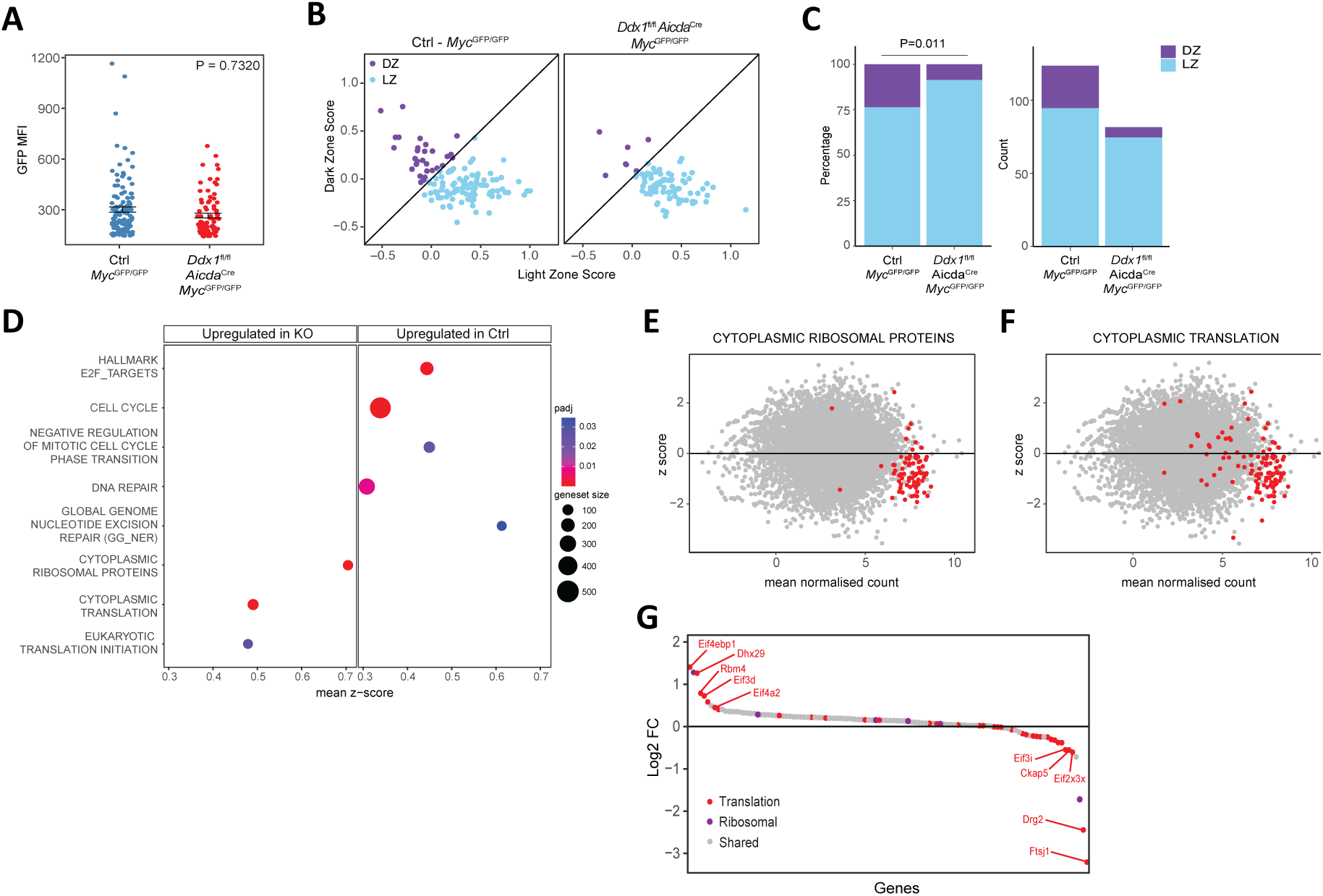
Single-cell RNA-Seq analyses of c-MYC^+^ germinal centre B-cells from *Ddx1*^fl/fl^ *Aicda*^Cre^ mice. Splenic MYC-eGFP^+^ GC B-cells were single-cell sorted from Ctrl *Myc*^GFP/GFP^ and *Ddx1*^fl/fl^ *Aicda*^Cre^ *Myc*^GFP/GFP^ mice, 10d post NP-KLH immunisation. **a**, Quantification of GFP MFI index-sorted as single MYC-eGFP^+^ GC B cells. Each data-point represents an individual cell, bar represents mean with standard error (n=204). The *P* value was generated by performing an unpaired, two-sided Student t-test on logarithmically transformed data. **b**, Dot plot showing DZ and LZ scores for each individual cell. Scores were assigned based on expression of transcripts described in Victora et al., 2012 and cells were attributed the phenotype associated with the highest score. Diagonal line represents the intercept between scores. **c**, Quantification of the frequency (left) and total number (right) of cells assigned to DZ and LZ in Ctrl *Myc*^GFP/GFP^ and *Ddx1*^fl/fl^ *Aicda*^Cre^ *Myc*^GFP/GFP^ mice. The *P* value was generated by performing a Pearson’s Chi-squared test. **d**, GSEA of differential gene expression between MYC-eGFP^+^ GC B-cells from Ctrl *Myc*^GFP/GFP^ and *Ddx1*^fl/fl^ *Aicda*^Cre^ *Myc*^GFP/GFP^ mice. Plot displays representative pathways that were identified as the most significant and non-overlapping gene-sets, stratified on those upregulated in *Ddx1*^fl/fl^ *Aicda*^Cre^ MYC-eGFP^+^ GC B-cells (left-hand side) and those upregulated in Ctrl MYC-eGFP^+^ GC B-cells (right-hand side). Size of points illustrates size of gene-set. The *P* values were generated by performing aKolmogorov–Smirnov test with Holm-Sidak correction for multiple testing. **e-f**, MA plots comparing gene expression in MYC-eGFP^+^ GC B-cells, with Z-scores <0 and >0 representing genes increased in Ctrl *Myc*^GFP/GFP^ and *Ddx1*^fl/fl^ *Aicda*^Cre^ *Myc*^GFP/GFP^ mice, respectively. Each grey point represents a gene identified in both datasets. Cytoplasmic ribosomal proteins (**e**) and cytoplasmic translation (**f**) gene-sets are shown. **g**, Log2 fold change (Log2FC) of transcript expression calculated for *Ddx1*^fl/fl^ *Aicda*^Cre^ MYC-eGFP^+^ GC B-cells over Ctrl MYC-eGFP^+^ GC B-cells. Grey-points represent transcripts shared between ribosomal protein and translation gene-sets. Purple and red points represent transcripts unique to ribosomal protein or translation gene-sets, respectively.

We determined the stage of the cell-cycle for MYC-eGFP^+^ GC B-cells based on gene expression profiles using the CellCycleScoring function within the Seurat R-package^38^. We detected cells in all three cell-cycle phases in both Ctrl *Myc*^GFP/GFP^ and *Ddx1*^fl/fl^ *Aicda*^Cre^ *Myc*^GFP/GFP^ mice (Supplementary Fig. 5, C and D), with a predominance for DZ B-cells to populate S/G2/M phases of the cell cycle (Supplementary Fig. 5E). Previous studies have shown that although LZ B-cells are predominantly in G1, a proportion undergoes cyclic re-entry prior to traversing back into the DZ, and these can complete one full cell division^6, 10^. In line with these observations, the majority of the isolated LZ GC B-cells in this analysis were in G1 phase of the cell-cycle (Supplementary Fig. 5F). In *Ddx1*^fl/fl^ *Aicda*^Cre^ *Myc*^GFP/GFP^ mice we observed a reduced proportion of LZ B-cells in the S-phase compared to Ctrl *Myc*^GFP/GFP^ mice, but this did not reach significance possibly due to the reduced number of LZ cells in this cell-cycle phase (Supplementary Fig. 5F). This demonstrates that cell-cycle scoring based on gene expression profiles recapitulates the known biology of GC B-cells, although we were unable to detect differences in cell-cycle phase distribution between MYC-eGFP^+^ GC B-cells from Ctrl *Myc*^GFP/GFP^ and *Ddx1*^fl/fl^ *Aicda*^Cre^ *Myc*^GFP/GFP^ mice.

To visualise differences between the gene expression profiles of Ctrl and DDX1-deficient MYC-eGFP^+^ GC B-cells we performed dimensionality reduction utilising t-distributed Stochastic Neighbour Embedding (t-SNE)^39^. There was no differential clustering between Ctrl *Myc*^GFP/GFP^ and *Ddx1*^fl/fl^ *Aicda*^Cre^ *Myc*^GFP/GFP^ mice (Supplementary Fig. 5G). Using a pseudo-bulk approach to examine the dataset further, we observed minimal gene expression changes with only two differentially expressed genes (DEGs) below the 0.05 significance threshold as determined by DESeq2^40^ (Supplementary Fig. 5H). To determine whether any gene pathways are differentially expressed between MYC-eGFP^+^ GC B-cells from Ctrl *Myc*^GFP/GFP^ and *Ddx1*^fl/fl^ *Aicda*^Cre^ *Myc*^GFP/GFP^ mice, Gene Set Enrichment Analysis (GSEA) was applied to the pseudobulk dataset using gene-sets described by Merico et al., 2010^41^. Pathways upregulated in MYC-eGFP^+^ GC B-cells from Ctrl *Myc*^GFP/GFP^ mice were associated with cell-cycle and DNA repair (Fig. 5D, Supp Table 3). Strikingly, the GSEA analysis in MYC-eGFP^+^ GC B-cells from *Ddx1*^fl/fl^ *Aicda*^Cre^ *My*c^GFP/GFP^ mice showed enrichment for pathways involved in cytoplasmic translation and ribosomal proteins (Fig. 5D, Supplementary Table 3). To identify the distribution of transcripts within these gene-sets, MA plots were generated and transcripts for cytoplasmic ribosomal proteins and translation gene-sets overlayed. This highlighted that most transcripts from both gene-sets had negative Z-scores in DDX1-deficient MYC-eGFP^+^ GC B-cells (Fig. 5, E and F). To identify those genes with the highest differential expression between genotypes, log2 fold-change (log2FC) was calculated for each transcript and plotted from increasing to decreasing score, with the occurrence among the translation and ribosomal gene-sets annotated (Fig. 5G, Supplementary Table 4). Transcripts with the highest and lowest log2FC originated from the cytoplasmic translation gene-set. These transcripts encode for subunits of the Eif4, Eif3 and Eif2 complexes and the RNA helicase Dhx29, all of which are involved in translation initiation. Interestingly, the most downregulated gene in DDX1-deficient MYC-eGFP^+^ GC B-cells encodes for the RNA 2’-O-Methyltransferase Ftsj1, which modifies tRNAs at positions 32 and 34 within the anticodon loop, thus determining translation efficiency^42^. Based on these findings we postulated that DDX1-deficient MYC-eGFP^+^ GC B-cells were transcriptionally poised to transition to the DZ but were unable to do so due to defects in the translation of these transcripts.

### DDX1 depletion impairs mRNA translation due to defects in tRNA splicing

To determine whether mRNA translation defects underlie the impaired proliferation of DDX1-deficient GC B-cells undergoing positive selection, we administered puromycin to Ctrl *My*c^GFP/GFP^ and *Ddx1*^fl/fl^ *Aicda*^Cre^ *My*c^GFP/GFP^ mice 10d post-immunisation and measured its incorporation into cells using flow cytometry. Puromycin is incorporated into elongating peptide chains by the ribosome and prevents further elongation, thus capturing ribosomes in the act of translation. The reduced puromycin incorporation by naïve B-cells compared to GC B-cells is consistent with their low protein biosynthetic activity^43^ (Supplementary Fig. 6, A and B). Amongst GC B-cells, the MYC-eGFP^+^ subset incorporated the greatest amount of puromycin (Supplementary Fig. 6, A and B), consistent with protein synthesis being induced by positive selection. We detected a significant reduction in puromycin incorporation in GC B-cells from *Ddx1*^fl/fl^ *Aicda*^Cre^ *My*c^GFP/GFP^ mice (Fig. 6A). Notably, while puromycin labelling intensity was highest in MYC-eGFP^+^ LZ B-cells, protein synthesis capacity was strongly impaired in this population in *Ddx1*^fl/fl^ *Aicda*^Cre^ *My*c^GFP/GFP^ mice (Fig. 6A). These results demonstrate that DDX1 is required for the boost in mRNA translation activity characteristic of positively selected GC B-cells.

**Figure 6.**
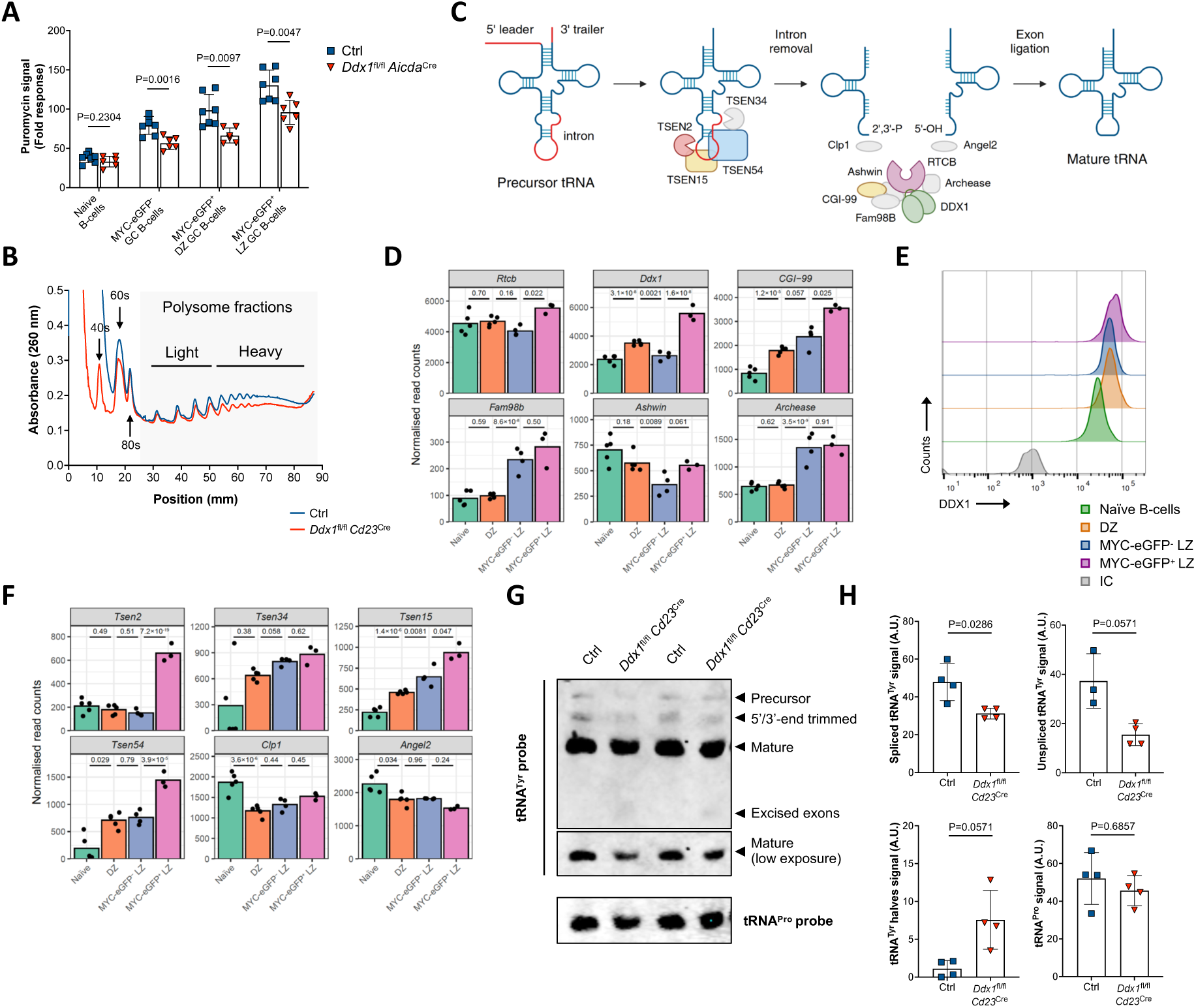
DDX1-deficient GC B-cells are impaired in mRNA translation due to defects in tRNA splicing. **a**, Quantification of puromycin signal measured by flow cytometry in naïve B-cells (CD19^+^IgD^+^) and GC B-cells from Ctrl *Myc*^GFP/GFP^ and *Ddx1*^fl/fl^ *Aicda*^Cre^ *Myc*^GFP/GFP^ mice, 10d post NP-KLH immunisation. GC B-cells were further subdivided in MYC-eGFP^-^ DZ, MYC-eGFP^+^ DZ and MYC-eGFP^+^ LZ subsets. Fold response for puromycin signal in different subsets is normalised to response in negative control mice for puromycin labelling (i.e. PBS injection). Each datapoint represents an individual mouse, bar represent mean ± SD. Data is from 2 independent experiments. The *P* values were generated by performing an unpaired, two-sided Student’s t-test on logarithmically transformed data. **b**, Sucrose gradient fractionation of B-cell lysates. Splenic B-cell cultures from Ctrl and *Ddx1*^fl/fl^ *Cd23*^Cre^ mice were analysed 17 hours post-stimulation with anti-IgM and IL-4. Data is representative of 3 independent experiments, each using a pool of 2 biological replicates for each genotype. **c**, Schematic diagram of the tRNA splicing pathway showing a multi-step process that requires several tRNA processing factors. Excision of the intron is carried out by the tRNA splicing endonuclease complex (TSEN) composed of two catalytic subunits (TSEN2 and TSEN34) and two structural subunits (TSEN54 an TSEN15). The cleavage reaction generates a 5’-exon containing a 2’3’-cyclic phosphate and a 3’-exon containing a 5’-OH; Clp1 and Angel2 modify tRNA 5’-exon and 3’-exon ends, respectively, preventing RNA ligation catalysed by RTCB. The tRNA ligase complex is composed of core subunits RTCB, DDX1, Ashwin, CGI-99 and Fam98B, and a protein co-factor known as Archease. tRNA processing factors which expression is modulated in positively selected GC B-cells are shown in colour. **d**, Expression levels of transcripts encoding tRNA-LC subunits in naïve B-cells, DZ, MYC-eGFP^-^ and MYC-eGFP^+^ LZ GC B-cell subsets. Cells were isolated from wild-type *Myc*^GFP/GFP^ mice immunised with SRBC for 8 days and used for bulk RNA-Seq. The *P* values were derived from pairwise DESeq2 analyses using a Wald test, with FDR correction for multiple testing. **e**, Representative histograms for intracellular DDX1 protein levels measured by flow cytometry in naïve B-cells, DZ, MYC-eGFP^-^ LZ and MYC-eGFP^+^ LZ B-cells from Ctrl *Myc*^GFP/GFP^ mice, 10d post NP-KLH immunisation. Signal obtained using an anti-DDX1 monoclonal antibody is compared to response for isotype control (IC). **f**, Expression levels of transcripts encoding TSEN complex subunits, Clp1 and Angel2, obtained from the same bulk RNA-Seq datasets as in (**d**). **g**, Northern blot analysis of RNA isolated from splenic B-cell cultures from Ctrl and *Ddx1*^fl/fl^ *Cd23*^Cre^ mice, analysed 19 hours post-stimulation with anti-IgM and IL-4. Data shown for 2 biological replicates for each genotype. **h**, Quantification of the abundance of the different tRNA species shown in (**e**), with values shown as arbitrary units (A.U.). Each datapoint represents B-cell cultures from an individual mouse, bar represent mean ± SD. Data is from 2 independent experiments. The *P* values were generated by performing an unpaired, two-sided Student’s t-test on logarithmically transformed data.

To determine the mechanism by which DDX1 affects mRNA translation, we used *ex vivo* stimulation of splenic B-cells to model high rates of protein synthesis activity associated with B-cell activation^18, 43^. Using puromycin labelling, we detected a sustained three-fold increase in the translation rate of B-cells cultured with anti-IgM and IL4 between 16-24h post-stimulation (Supplementary Fig. 6C). At the peak of protein synthesis, *Ddx1*^fl/fl^ *Cd23*^Cre^ B-cell cultures showed reduced puromycin incorporation relative to Ctrl (Supplementary Fig. 6D). This defect was not due to impaired activation of B-cells isolated from *Ddx1*^fl/fl^ *Cd23*^Cre^ mice as the induced expression of activation markers CD69 and CD86 was comparable to Ctrl (Supplementary Fig. 6, E and F). To further characterise how DDX1 affects mRNA translation, we profiled mRNAs based on the level of their association with ribosomes using sucrose gradient fractionation. We observed a reduced proportion of polysome-bound mRNAs in B-cells from *Ddx1*^fl/fl^ *Cd23*^Cre^ mice compared to Ctrl mice (Fig. 6B). Notably, this reduction was mostly evident for heavy polysomes, which contain mRNAs with the highest ribosome content. These results indicate that loss of DDX1 specifically impairs mRNA translation efficiency, at a time when B-cell activation is established, and B-cells are preparing for rapid rounds of proliferation (Supplementary Fig. 6G).

DDX1 was shown to promote the activity of the tRNA-LC, facilitating the joining of two tRNA exons by the RNA ligase RTCB after tRNA splicing^20^. We hypothesised that diminished mRNA translation in DDX1-deficient GC B-cells was due to defects in the processing of intron-containing tRNAs (Fig. 6C). To evaluate whether positive selection was associated with changes in the expression of tRNA processing factors, we mined a publicly available RNA-Seq dataset for naïve and GC B-cell subsets. Alongside RTCB and DDX1, we determined the expression of three other tRNA-LC core subunits, CGI-99, FAM98B and Ashwin^44^. Positive selection in the GC was accompanied by upregulation of transcripts encoding for RTCB, DDX1 and CGI-99, all of which are required for complex integrity (Fig. 6D). Furthermore, we detected increased levels of DDX1 protein in MYC-eGFP^+^ LZ B-cells compared to other GC B-cell subsets and naïve B-cells (Fig. 6E). While FAM98B (also part of the tRNA-LC core) and the tRNA-LC protein co-factor Archease were upregulated in LZ GC B-cells, Ashwin expression was variable amongst subsets (Fig. 6D). Further corroborating the requirement for tRNA-LC activity during the GC response, we detected a 3-fold increase in the expression of transcripts encoding for the tRNA splicing endonuclease TSEN2, as well as upregulation of two structural subunits of the TSEN complex, TSEN15 and TSEN54 (Fig. 6F). Interestingly, expression of the 5’-hydroxyl polynucleotide kinase Clp1 and the 2’3’-cyclic phosphatase Angel2 (both known to counteract the activity of the tRNA-LC through modification of tRNA exon ends) showed the opposite trend, with decreased mRNA levels detected in GC B-cell subsets compared to naïve B-cells (Fig. 6F). Taken together, these results demonstrate that positive selection in the GC is supported by increased expression of various tRNA processing factors along the tRNA splicing pathway.

To determine whether DDX1 deficiency results in weakened activity of the tRNA-LC, we evaluated the level of tRNAs decoding tyrosine amino acid (tRNA^Tyr^) in *ex vivo* stimulated B-cells from Ctrl and *Ddx1*^fl/fl^ *Cd23*^Cre^ mice. Splicing of tRNA^Tyr^ provides a direct measure of tRNA-LC activity as all 10 genes encoding the tRNA^Tyr^ family contain an intron. We observed a decrease in the amount of precursor and spliced tRNA^Tyr^, and the presence of tRNA^Tyr^ exon halves, in B-cells from *Ddx1*^fl/fl^ *Cd23*^Cre^ mice (Fig. 6G and H). Nevertheless, a substantial amount of spliced tRNA^Tyr^ was still present in *Ddx1*^fl/fl^ *Cd23*^Cre^ B-cells, likely due to high stability of tRNA species, and the non-proliferative nature of B-cells within the first 24h of anti-IgM and IL-4 stimulation (Fig. 6G). To control for total RNA loading, we probed for splicing-independent tRNA^Pro^ and detected comparable expression levels in B-cells from *Ddx1*^fl/fl^ *Cd23*^Cre^ and Ctrl mice (Fig. 6G and H). This demonstrates the requirement of DDX1 for efficient splicing of tRNA^Tyr^ and suggests a more general role for DDX1 in modulating the abundance of intron-containing tRNA molecules.

Overall, our findings indicate that an increase in the kinetics of protein translation primes B-cells for rapid cell proliferation. In the absence of DDX1, decreased tRNA-LC activity and incomplete processing of intron-containing tRNAs, renders elongating ribosomes sensitive to changes in the availability of functional tRNAs.

## Discussion

Clonal expansion of GC B-cells with enhanced antigen-affinity is a core requisite for effective humoral immunity. Here, we show that ablation of the RNA helicase DDX1 renders GC B-cells unable to undergo rapid rounds of cell division in response to T_FH_-mediated selection and upregulation of the transcription factor c-MYC. DDX1 is required for tRNA-LC activity and splicing of intron-containing tRNAs, thus determining the extent of mRNA translation capacity following positive selection in GC B-cells. These studies establish tRNA maturation and the efficiency of protein synthesis as a new determinant of GC B-cell fitness during the proliferative burst that expands antigen-specific, high-affinity B-cell clones.

We demonstrate DDX1’s function during GC maturation is independent of the AID targeting activity we have previously described for DDX1 in CSR^19^. Previous studies show that CSR occurs prior to GC formation at the T:B border^45, 46, 47, 48^, with the increase in IgG1^+^ GC B-cells mediated through positive selection of class-switched B-cells as the GC response matures^49^. Our findings align with these observations, as we detected no impact on the proportion of class-switched IgG1^+^ GC B-cells at 7d post-immunisation, most likely due to *Aicda*^Cre^-mediated DDX1 protein depletion occurring after CSR has taken place. Analysis of nucleotide substitutions in the J_H_4 intron indicates SHM is not impaired in *Ddx1*^fl/fl^ *Aicda*^Cre^ mice. However, our studies do not formally exclude a role for DDX1 in SHM due to incomplete protein depletion at the early stages of the GC response in this conditional *Ddx1* knock-out model. Our finding that DDX1-deficiency impairs GC maturation in an AID-deficient background provides unequivocal evidence that DDX1 function during positive selection in the GC is uncoupled from its AID targeting activity.

DDX1-depleted GC B-cells were inherently impaired in their ability to clonally expand despite normal T_FH_-mediated upregulation of c-MYC and the establishment of the transcriptional programme characteristic of a positively selected GC B-cell. At a functional level, lack of DDX1 activity resulted in an almost complete absence of antigen-specific GC B-cells and affinity maturation of antibody responses. Recently it has been shown that T_FH_-cell mediated selection is not required for LZ GC B-cells to initiate cyclic re-entry, but the strength of T-cell help is essential to ‘refuel’ GC B-cells for subsequent rounds of division in the DZ^10^. The nature of this ‘refuelling’ mechanism is only partially understood but both the amount of MYC protein expressed in LZ B-cells and subsequent induction of CyclinD3 expression seem to determine the number of cycles a GC B-cell undergoes in the DZ^9, 12^. Together with mTORC1 signalling, MYC promotes the anabolic cell growth that sustains cell division in the DZ^11^, which likely results in increased ribosome biogenesis and overall protein biosynthetic capacity^50, 51, 52^. In the absence of DDX1, GC B-cells have reduced ability for protein synthesis which is manifested by the absence of highly translating c-MYC^+^ GC B-cells in the LZ and defects in mRNA translation upon B-cell activation. Thus, we propose that DDX1’s control over tRNA maturation is required to fulfil the biosynthetic impetus endowed to GC B-cells upon positive selection, to meet the increase in mRNA translation activity in response to c-MYC signalling.

In addition to tRNA splicing, the activity of the RTCB is required for expression of the spliced isoform of the transcription factor XBP1 (XBP1s). XBP1s is critical during PC differentiation, maintaining endoplasmic reticulum homeostasis as part of the unfolded protein response (UPR), thus enabling the generation and secretion of large quantities of antibodies^53, 54, 55^. Disruptions in RTCB expression prevents XBP1s upregulation in PC, resulting in decreased antibody secretion^21^. It is currently unknown whether DDX1 is required for XBP1s expression during PC differentiation. We note that XBP1 expression is not required for GC responses^56^ and we find that *Ddx1*^fl/fl^ *Aicda*^Cre^ mice express normal levels of antigen-specific IgM antibodies in the serum upon immunisation. Therefore, our findings indicate that the requirement for DDX1 in GC B-cells specifically relates to RTCB activity in the maturation of intron-containing tRNAs. Furthermore, RTCB-deficient plasmablasts were shown to have decreased levels of splicing-dependent tRNAs but this did not result in global changes in protein synthesis rates^21^. We reason this reflects distinct requirements for protein synthesis efficiency in GC B-cells and plasmablasts.

Modulation of tRNA expression has been shown to associate with cell proliferation and tumour progression^57, 58, 59, 60^. More specifically, the upregulation of intron-containing tRNA^Leu^ and tRNA^Tyr^ expression via mTORC signalling is coupled with the escape from senescence in tumour cells^61^. Also, depletion of tRNA^Tyr^ during oxidative stress led to impaired translation of growth and metabolic genes enriched in tyrosine codons^62^. It is possible that tRNA splicing is specifically required as GC B-cells transition from the LZ to the DZ, to change tRNA availability and selectively enhance protein synthesis based on differential codon usage. However, recent studies suggest that despite extensive tRNA repertoire remodelling during differentiation, tRNA anticodon pools remains largely stable^63^. Similarly, a translation boost of mRNAs containing rare codons underpins cell proliferation, but this is achieved by an overall increase in tRNA availability rather than differences in the expression of individual tRNAs^64^. This suggests that mechanisms other than relative tRNA abundance are likely to play a key role in calibrating gene expression.

Post-transcriptional, enzyme-catalysed modifications to tRNAs impact codon-anticodon recognition and regulate translational efficiency. In antibody-secreting PC, high rates of antibody production rely on tRNA adenine-to-inosine wobble modification, to meet biased codon demands of immunoglobulin genes^65^. Also, *N*^1^-methyladenosine tRNA modification at position 58 is associated with T-cell activation by controlling the efficiency of protein synthesis for c-MYC and other key proteins^66^. Notably, tRNA introns can serve as important substrate recognition elements for tRNA modifying enzymes. In *Saccharomyces cerevisiae*, the presence of an intron in tRNA^Tyr^ and tRNA^Leu^ is required for modifications of U35 to Ψ35 and C34 to m^5^C34, respectively, which impact on the efficiency of codon recognition^67, 68, 69^. Thus, impaired tRNA splicing in GC B-cells may alter further tRNA remodelling leading to changes in tRNA functionality during mRNA translation. Interestingly, we observed downregulation of the tRNA 2’-O-Methyltransferase Ftjs1 in positively selected GC B-cells upon DDX1 depletion. Although it is currently unclear whether Ftsj1-mediated modification of tRNA’s anticodon loop is linked to tRNA splicing, this result suggests that DDX1-deficiency can impact tRNA maturation beyond its immediate role in promoting tRNA-LC activity.

Our study shows that GC B-cell selection and affinity maturation of the antibody response are particularly sensitive to changes in the pool of functional tRNAs that B-cells have available. Understanding how B-cells utilise the vast regulatory potential of tRNAs to increase mRNA translation activity and achieve rapid clonal expansion is a new avenue for investigation. It is likely that modulation of tRNA levels and/or tRNA modification can be leveraged in the future for therapeutic intervention in antibody-mediated disease.

## Acknowledgments

We thank the Biological Support Unit (BSU03), Flow Cytometry Facility (Flowcyt04), Imaging Facility (Imag07) and the Bioinformatics Group (Bioinf01) at the Babraham Institute for technical assistance; these facilities receive financial support from the Babraham Institute’s BBSRC Core Capability Grant (CCG) BB/CCG2310/1. We thank Iain Macaulay and the Genomics Team at the Earlham Institute for single-cell RNA-seq analyses. We thank Martin Turner for his support and advice at different stages of this project; Martin Turner, Anne Corcoran, Kai-Michael Toellner and Alexander Saveliev for providing critical feedback on the manuscript; Jon Houseley and Danny Nedialkova for their guidance with northern blot analyses. This work was supported by a Wellcome Trust Henry Dale Fellowship (220196/Z/20/Z) to C.R.A and funding from BBSRC Institute Strategic Programmes Grant (ISPG) for Immunology BB/Y006917/1 and BBSRC Institute Development Grant BB/IDG2310/1. R.K. was supported by a BBSRC DTP studentship.

## Author contributions

R.K. conducted most of the experiments with help from F.S. for influenza infections, A.W. for northern blot analyses, J.A. and D.A. for VDJ sequencing and M.S. for J_H_4 sequencing. C.R.A. performed polysome fractionation experiments with help from M.S. S.I. performed bioinformatics analyses of scRNA-seq data, supervised by S.A. H.O. developed methodology for immunofluorescence image processing and quantification. R.K. and C.R.A conceptualized the study and wrote the manuscript. C.R.A. supervised the study and acquired funding.

## Supplementary Figures

**Supplementary Figure 1:**
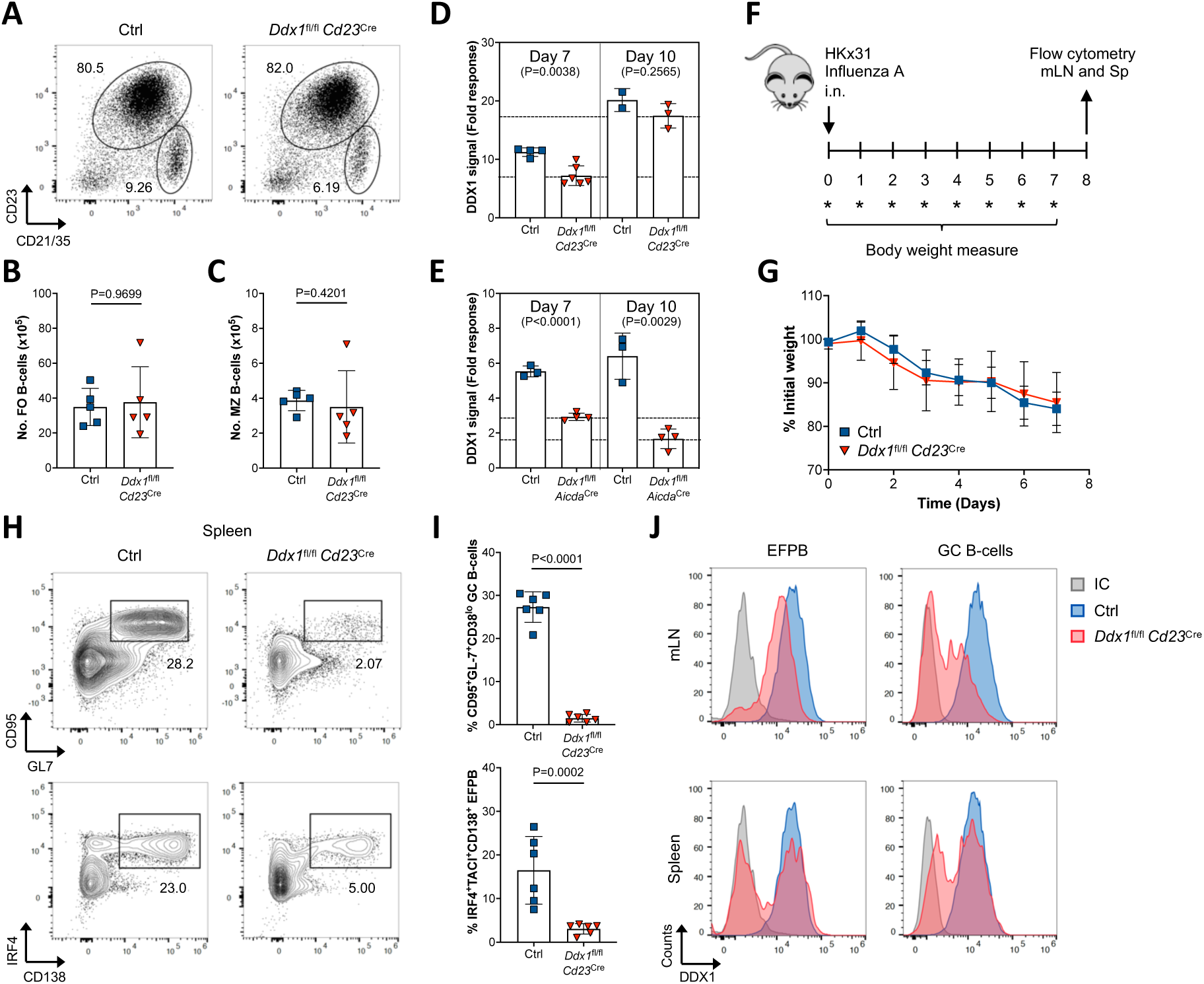
(**a**) Representative flow cytometry plots of FO (CD23^+^CD21/35^lo^) and MZ (CD23^lo^CD21/35^+^) CD19^+^B220^+^ B-cells, in spleens from Ctrl and *Ddx1*^fl/fl^ *Cd23*^Cre^ mice. (**b,c**) Quantification of the total numbers of FO (**b**) and MZ (**c**) B-cells gated in (**a**). Each datapoint represents an individual mouse, bar represent mean ± SD. Data is from 6 independent experiments. The *P* values were generated by performing an unpaired, two-sided Student’s t-test on logarithmically transformed data. (**d,e**) Quantification of geometric mean fluorescent intensity (geoMFI) for intracellular DDX1 protein levels measured by flow cytometry in GC B-cells from *Ddx1*^fl/fl^ *Cd23*^Cre^ (**d**), *Ddx1*^fl/fl^ *Aicda*^Cre^ (**e**) and Ctrl mice, 7d and 10d post-immunisation with NP-KLH. Dashed lines represent average values for DDX1 cKO mice at d7 and d10. Data normalised to response for isotype control (IC) (Fold response). Each datapoint represents an individual mouse, bars represent mean ± S.D. Data is from 2-3 experiments. The *P* values were generated by performing an unpaired, two-sided Student’s t-test on logarithmically transformed data. (**f**) Experimental approach in which Ctrl and *Ddx1*^fl/fl^ *Cd23*^Cre^ mice were intranasally (i.n.) inoculated with 10^4^ CFU of HKx31 Influenza A. Mice were taken down 8d post-infections and mediastinal lymph nodes (mLN) and spleens (Sp) were isolated and processed for flow cytometric analysis. (**g**) Quantification of weight in HKx31 Influenza A infected animals. Weight loss is represented as percentage decrease from initial body weight measurement. Each datapoint represents mean ± SD for each group, with 4-6 mice per group. (**h**) Representative flow cytometry plots of GC B-cells (CD95^+^GL-7^+^) in CD138^-^IgD^-^CD19^+^ B-cells (top) and EFPB (IRF4^+^CD138^+^) in IgD^-^CD19^+^ B-cells (bottom) in spleens from Ctrl and *Ddx1*^fl/fl^ *Cd23*^Cre^ mice, 8d post HKx31 Influenza A infection. (**i**) Quantification of the frequency of GC B-cells (top) and EFPB (bottom) defined as CD95^+^GL-7^+^CD38^lo^ and IRF4^+^CD138^+^TACI^+^, respectively. Each datapoint represents an individual mouse, bar represent mean ± SD. Data is from 2 independent experiments. The *P* values were generated by performing an unpaired, two-sided Student’s t-test on logarithmically transformed data. (**j**) Representative histograms for intracellular DDX1 protein levels measured by flow cytometry in GC B-cells and EFPB in mediastinal lymph nodes (mLN) and spleens (Sp) from Ctrl and *Ddx1*^fl/fl^ *Cd23*^Cre^ mice, 8d post HKx31 Influenza A infection. Signal obtained using an anti-DDX1 monoclonal antibody is compared to response for isotype control (IC).

**Supplementary Figure 2:**
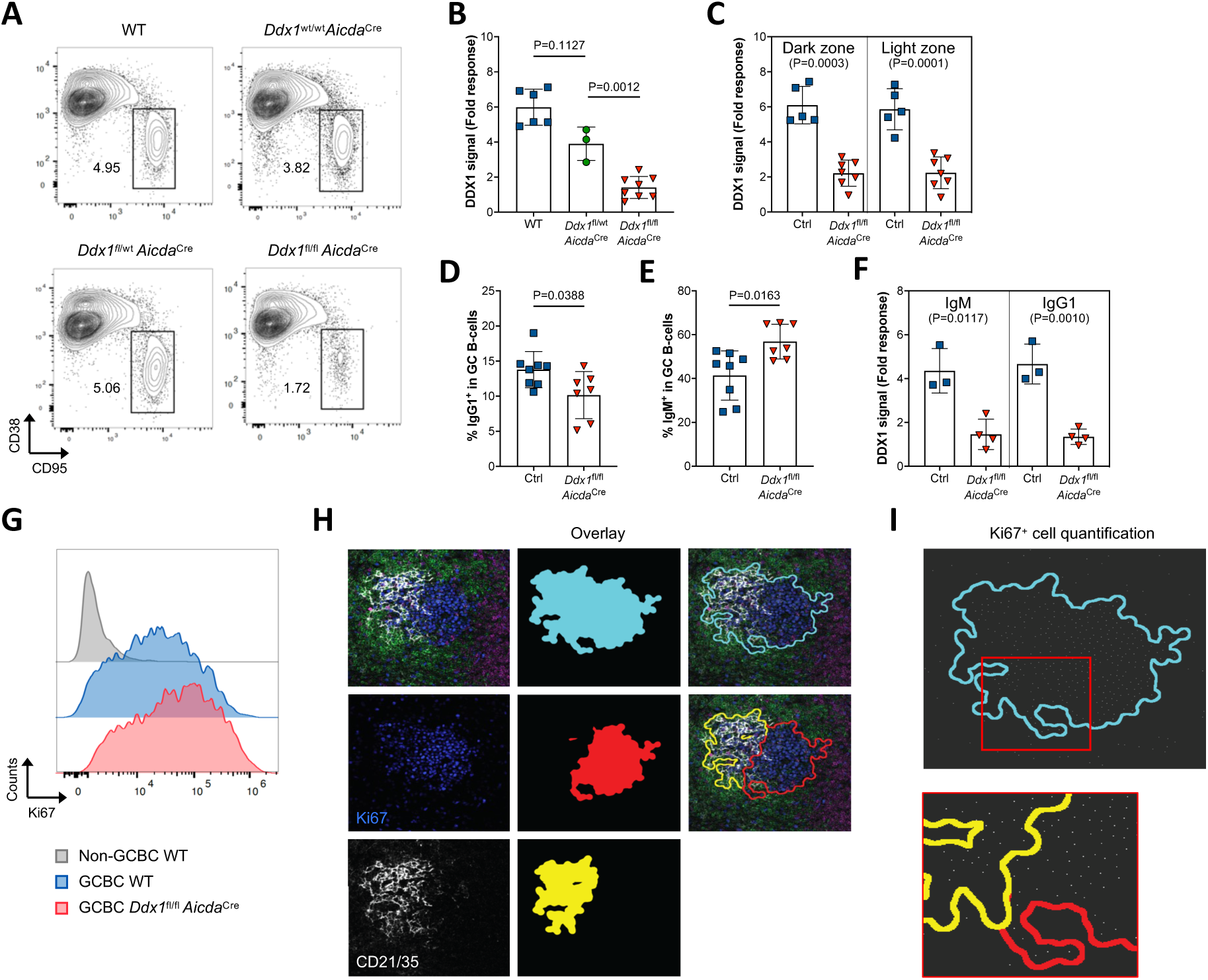
(**a**) Representative flow cytometry plots of GC B-cells (CD95^+^CD38^lo^) within the total CD19^+^B220^+^ B-cell population in spleens from wild-type (WT), *Ddx1*^wt/wt^ *Aicda*^Cre^, *Ddx1*^fl/wt^ *Aicda*^Cre^ and *Ddx1*^fl/fl^ *Aicda*^Cre^ mice, 10d post NP-KLH immunisation. (**b**) Quantification of geometric mean fluorescent intensity (geoMFI) for intracellular DDX1 protein levels measured by flow cytometry in GC B-cells. Data normalised to response for isotype control (IC) (Fold response). Each datapoint represents an individual mouse, bars represent mean ± S.D. Data is from 3 independent experiments. The *P* values were generated by performing a one-way ANOVA on logarithmically transformed data, with Holm-Sidak correction for multiple testing. (**c**) Quantification of geoMFI for intracellular DDX1 protein levels measured by flow cytometry in GC B-cells stratified as DZ/LZ. Each datapoint represents an individual mouse, bars represent mean ± S.D. Data is from 2 independent experiments. The *P* values were generated by performing an unpaired, two-sided Student’s t-test on logarithmically transformed data. (**d**,**e**) Quantification of percentage of IgG1^+^ (**d**) and IgM^+^ (**e**) GC B-cells in Ctrl and *Ddx1*^fl/fl^ *Aicda*^Cre^ mice. Each datapoint represents an individual mouse, bars represent mean ± S.D. Data is from 3 independent experiments. The *P* values were generated by performing an unpaired, two-sided Student’s t-test on logarithmically transformed data. (**f**) Quantification of geoMFI for intracellular DDX1 protein levels measured by flow cytometry in GC B-cells stratified as IgM^+^/IgG1^+^. Each datapoint represents an individual mouse, bars represent mean ± S.D. Data is from one experiment. The *P* values were generated by performing an unpaired, two-sided Student’s t-test on logarithmically transformed data. (**f**) Representative histograms of Ki67 expression measured by flow cytometry in GC B-cells (CD38^lo^CD95^+^CD19^+^B220^+^) from Ctrl and *Ddx1*^fl/fl^ *Aicda*^Cre^ mice in comparison to signal from non-GC B-cells (CD38^+^CD95^-^CD19^+^B220^+^). (**g**) Representative CellProfiler overlay to quantify total GC area, DZ and LZ areas. Cyan outline represents the automated GC structures identified, and DZ and LZ areas are represented by delimitation with red and yellow outlines, respectively. (**h**) Representative images used for quantification of the numbers of Ki67^+^ cells within GC, DZ and LZ areas using CellProfiler and ImageJ. Each white point is a Ki67^+^ cell defined by DAPI nuclear staining and intensity in Ki67 channel. Full image and zoomed in sections are depicted for visualisation.

**Supplementary Figure 3:**
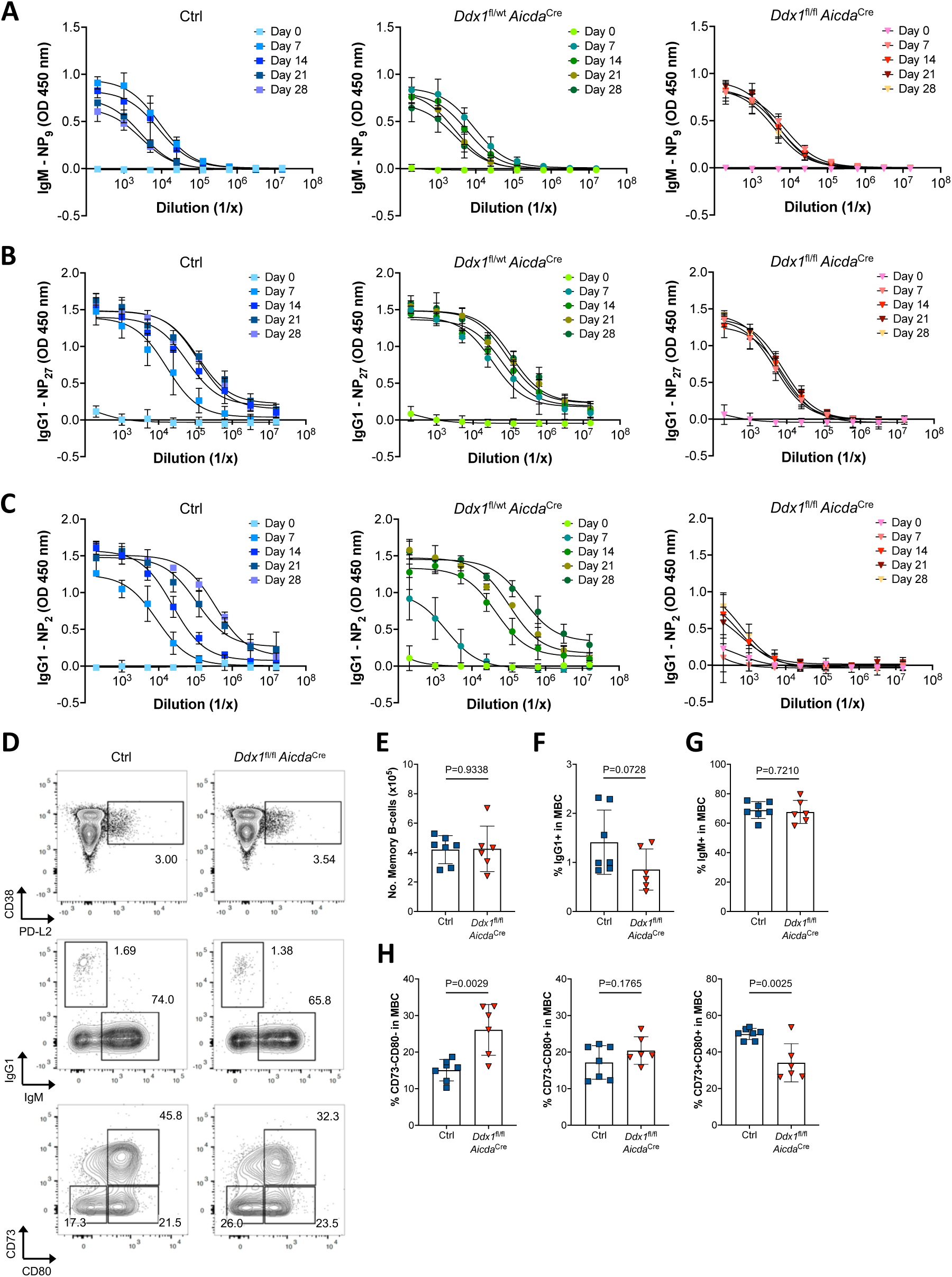
(**a-c**) Serial dilution curves for quantification of NP(9)-IgM antibodies (**a**), broad-affinity NP(27)-IgG1 antibodies (**b**) and high-affinity NP(2)-IgG1 antibodies (**c**) in the serum of Ctrl, *Ddx1*^fl/wt^ *Aicda*^Cre^ and *Ddx1*^fl/fl^ *Aicda*^Cre^ mice collected 0d, 7d, 14d, 21d and 28d post-immunisation with NP-KLH. Each datapoint represents the average optical density (OD) value at 450nm obtained for the respective serum dilution in 4/5 mice per group (mean ± S.D). (**d**) Representative flow cytometry plots of MBC (CD38^int^PD-L2^+^) within the CD19^+^IgD^-^CD95^-^ B-cell population in spleens from Ctrl and *Ddx1*^fl/fl^ *Aicda*^Cre^ mice, 10d post NP-KLH immunisation. MBC are further defined based on the expression of IgG1 and IgM, as well as CD80 and CD73. (**e-g**) Quantification of the total numbers of MBC (**e**) and the frequency of IgG1^+^ (**f**) and IgM^+^ MBC (**g**) gated in (**d**). (**h**) Quantification of the frequency of CD73^-^CD80^-^, CD73^-^CD80^+^ and CD73^+^CD80^+^ MBC subsets, gated in (**d**). Each datapoint represents an individual mouse, bar represent mean ± S.D (n=13). Data is from 2 independent experiments. The *P* values were generated by performing an unpaired, two-sided Student’s t-test on logarithmically transformed data.

**Supplementary Figure 4:**
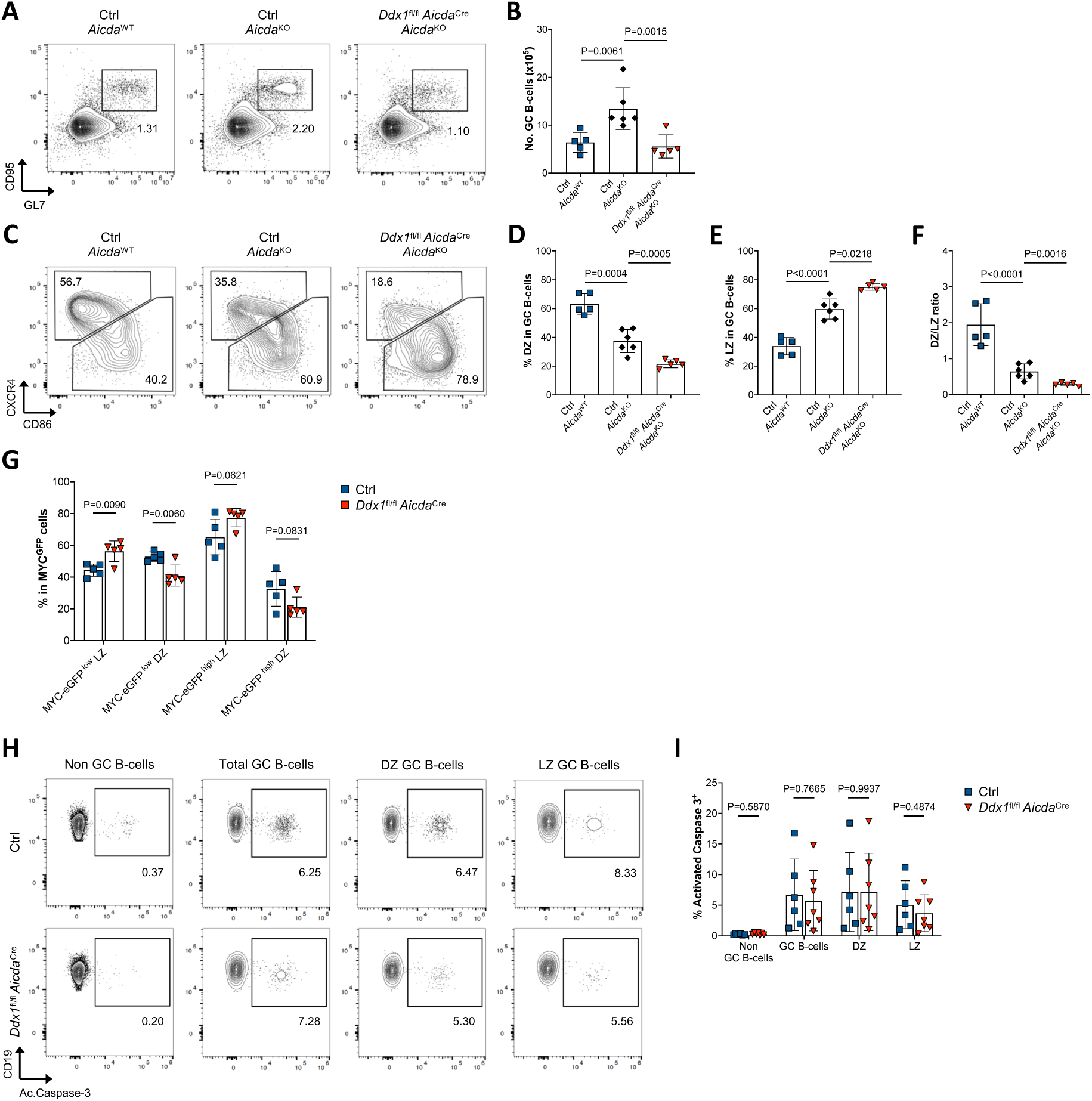
(**a**) Representative flow cytometry plots of GC B-cells (CD95^+^GL-7^+^) within total CD19^+^B220^+^ B-cell population in spleens from Ctrl *Aicda*^WT^, Ctrl *Aicda*^KO^ and *Ddx1*^fl/fl^ *Aicda*^Cre^ *Aicda*^KO^ mice, 10d post NP-KLH immunisation. (**b**) Quantification of total numbers of GC B-cells (defined as CD95^+^GL-7^+^PNA^+^). (**c**) Representative flow cytometry plots of DZ and LZ within GC B-cell population (CD38^lo^CD95^+^CD19^+^B220^+^). (**d-f**) Quantification of the percentage of DZ (**d**) and LZ (**e**) B-cells gated in (**c**), and DZ/LZ ratios (**f**). Each datapoint represents an individual mouse, bar represent mean ± S.D. Data is from 2 independent experiments. The *P* values were generated by performing a one-way ANOVA on logarithmically transformed data, with Holm-Sidak correction for multiple testing. (**g**) Quantification of the frequency of DZ and LZ subsets within MYC-eGFP^low^ and MYC-eGFP^high^ GC B-cell population. Each datapoint represents an individual mouse, bar represent mean ± SD. Data is from one experiment. The *P* values were generated by performing an unpaired, two-sided Student’s t-test on logarithmically transformed data. (**h**) Representative flow cytometry plots of activated caspase-3 expression in non-GC B-cells (CD38^+^CD95^-^CD19^+^), total GC, DZ and LZ B-cells. (**i**) Quantification of the frequency of activated caspase-3 in cell populations gated in (**h**). Each datapoint represents an individual mouse, bar represent mean ± S.D (n=13). Data is from 3 independent experiments. The *P* values were generated by performing an unpaired, two-sided Student’s t-test on logarithmically transformed data.

**Supplementary Figure 5:**
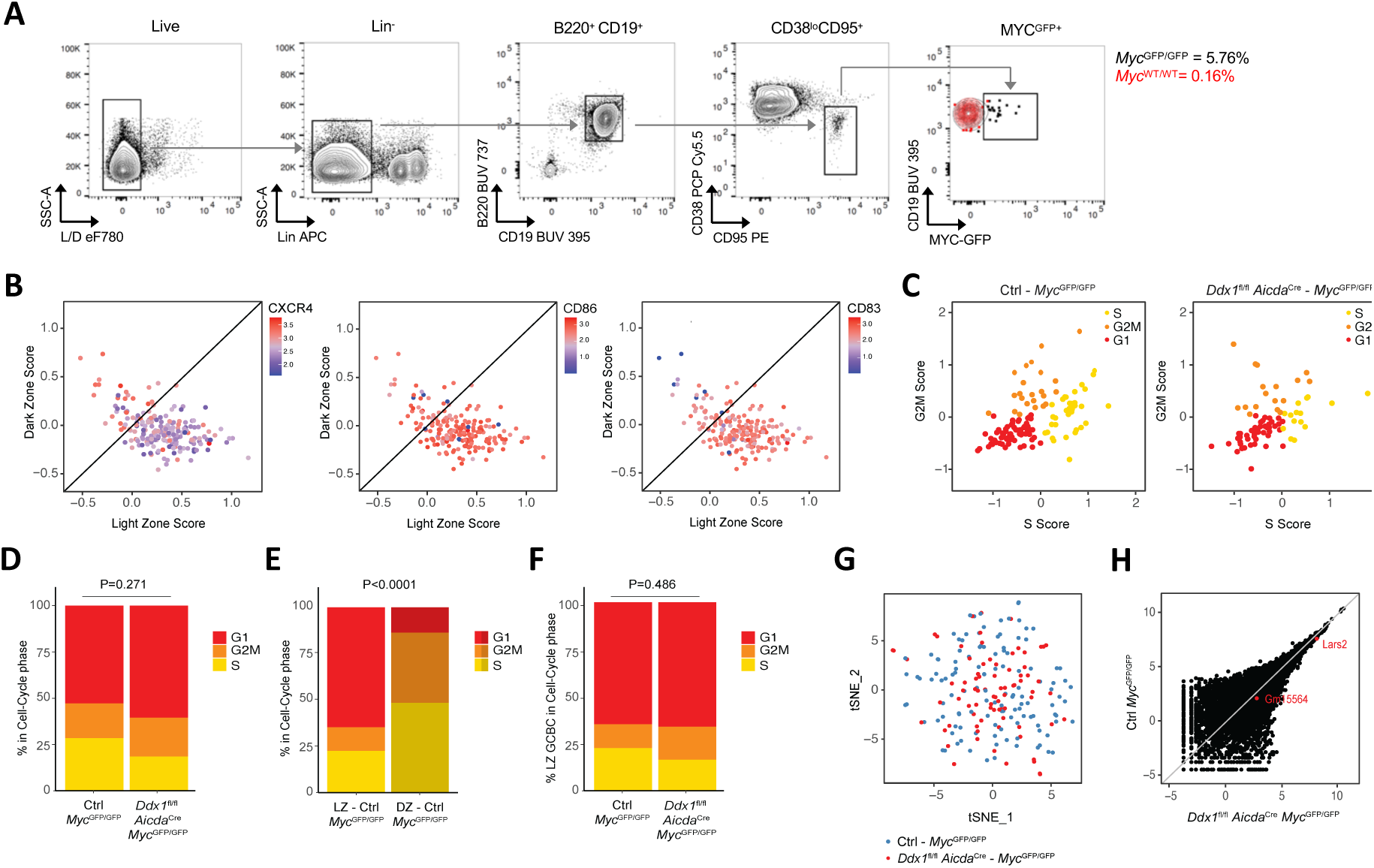
(**a**) Representative flow cytometry plots showing the gating strategy for single-cell flow sorting of splenic MYC^GFP+^ GC B-cells from Ctrl *Myc*^GFP/GFP^ and *Ddx1*^fl/fl^ *Aicda*^Cre^ *Myc*^GFP/GFP^ mice, 10d post NP-KLH immunisation. An overlay of MYC^GFP^ negative GC B-cells is shown in red. (**b**) Relative expression of CXCR4 (left), CD86 (middle) and CD83 (right) plotted on LZ against DZ score plots representing individual cells from Ctrl *Myc*^GFP/GFP^ and *Ddx1*^fl/fl^ *Aicda*^Cre^ *Myc*^GFP/GFP^ mice. (**c**) Individual cells from Ctrl *Myc*^GFP/GFP^ and *Ddx1*^fl/fl^ *Aicda*^Cre^ *Myc*^GFP/GFP^ mice were assigned to a cell-cycle phase dependant on transcript profiles. Cells were coloured based on assignment in S (yellow), G1 (red) and G2M (orange). (**d,e**) Quantification of the frequency of cells in each phase of the cell-cycle stratified by genotype (**d**) and LZ/DZ distribution in Ctrl *Myc*^GFP/GFP^ mice (n=123) (**e**). (**f**) Quantification of the frequency of cells in each phase of the cell-cycle stratified on LZ GC B-cells and genotype (n=168). The *P* value was generated by performing a Pearson’s Chi-squared test. (**g**) Dimensionality reduction utilising t-distributed Stochastic Neighbour Embedding (t-SNE); each dot represents a single-cell coloured based on genotype. (**h**) Pseudobulk analysis of differential gene expression between Ctrl *Myc*^GFP/GFP^ and *Ddx1*^fl/fl^ *Aicda*^Cre^ *Myc*^GFP/GFP^ mice. Each datapoint represents an individual gene; differentially expressed genes between Ctrl and *Ddx1*^fl/fl^ *Aicda*^Cre^ MYC^GFP+^ GC B-cells identified by DESeq2 function are highlighted in red.

**Supplementary Figure 6:**
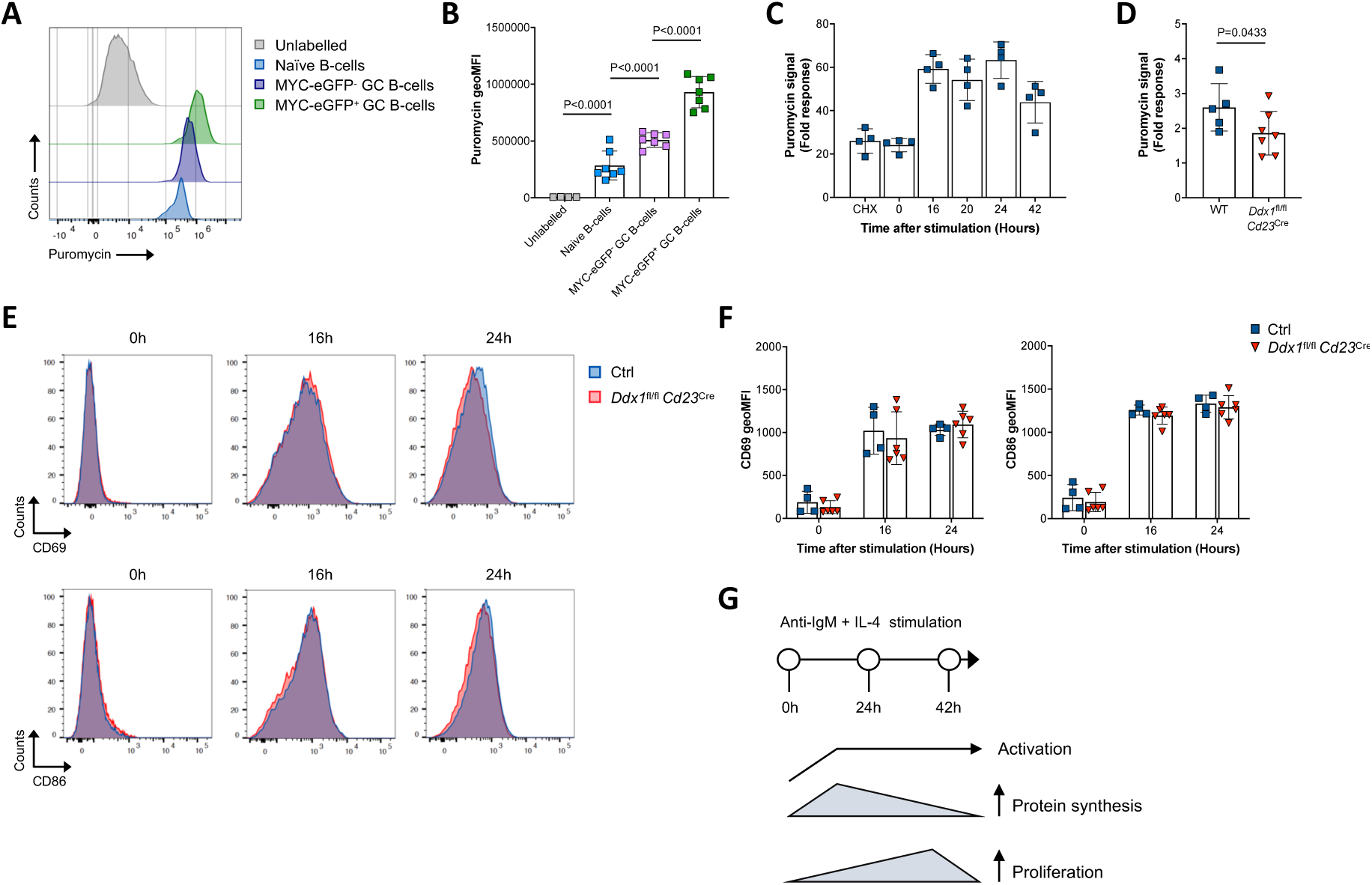
(**a**) Representative histograms of puromycin signal measured by flow cytometry in naïve B-cells (CD19^+^IgD^+^), MYC-eGFP^-^ and MYC-eGFP^+^ GC B-cells from Ctrl *Myc*^GFP/GFP^ mice, 10d post NP-KLH immunisation. Histogram for unlabelled cells demonstrates the signal in total GC B-cells from mice that received a I.P. injection of PBS at the time of puromycin administration. (**b**) Quantification of geoMFI for puromycin signal displayed in (**a**). Each data point represents an individual mouse, bar represent mean ± S.D. Data is from 2 independent experiments. The *P* values were generated by performing a one-way ANOVA on logarithmically transformed data, with Holm-Sidak correction for multiple testing. (**c**) Quantification of puromycin signal in splenic B-cell cultures from Ctrl mice, shown as fold response over PBS. B-cells were pulsed with puromycin for 15 minutes at 0, 16, 20 and 24 hours post-stimulation with anti-IgM and IL-4. Treatment with cycloheximide (CHX) 5 minutes prior to puromycin pulse was used as a negative control for actively translating ribosomes. (**d**) Quantification of puromycin signal in splenic B-cell cultures from Ctrl and *Ddx1*^fl/fl^ *Cd23*^Cre^ mice, analysed 16 hours post-stimulation with anti-IgM and IL-4. Data is shown as fold response over signal obtained for CHX-treated B-cells. Each datapoint represents B-cell cultures from an individual mouse, bar represent mean ± SD. Data is from 3 independent experiments. The *P* values were generated by performing an unpaired, one-sided Student’s t-test on logarithmically transformed data. (**e**) Representative histogram overlays of CD69 and CD86 signal measured by flow cytometry in splenic B-cell cultures from Ctrl and *Ddx1*^fl/fl^ *Cd23*^Cre^ mice, analysed 0, 16 and 24 hours post-stimulation with anti-IgM and IL-4. (**f**) Quantification of geoMFI for CD69 and CD86 signal displayed in (**e**). Each datapoint represents B-cell cultures from an individual mouse, bar represent mean ± SD (n=10). Data is from 2 independent experiments. The *P* values were generated by performing an unpaired, two-sided Student’s t-test on logarithmically transformed data. (**g**) Schematic diagram of B-cell activation following anti-IgM and IL-4 stimulation. A peak in protein synthesis efficiency is achieved at the time B-cell activation is established (i.e. 16h post stimulation). This precedes rapid rounds of cell proliferation, that are characteristic of lymphocyte activation.

## Methods

### Animal models

Mice were bred and maintained in the Babraham Institute Biological Support Unit (BSU). Mice were housed under pathogen-free conditions. No primary pathogens or additional agents listed in the FELASA recommendations have been confirmed during health monitoring surveys of the stock holding rooms. Ambient temperature was ∼19–21°C and relative humidity 52%. Lighting was provided on a 12-hour light: 12-hour dark cycle including 15 min ‘dawn’ and ‘dusk’ periods of subdued lighting. After weaning, mice were transferred to individually ventilated cages with 1-5 mice per cage. Mice were fed CRM (P) VP diet (Special Diet Services) ad libitum and received seeds (e.g., sunflower, millet) at the time of cage cleaning as part of their environmental enrichment. All mouse experimentation was approved by the Babraham Institute Animal Welfare and Ethical Review Body. Animal husbandry and experimentation complied with European Union and United Kingdom Home Office legislation. The following transgenic mice strains were used: *Ddx1^fl/fl^* (*Ddx1*^tm1c(EUCOMM)Hmgu^ allele) *Cd23*^Cre^ (described in Kwon et al, 2008^70^), *Aicda*^Cre^ (described in Kwon et al, 2008^70^), *Aicda*^KO^ (*Aicda*^tm1b(EUCOMM)Hmgu^ allele) and *Myc*^GFP^ (described in Huang et al, 2008^30^); with all transgenic mice maintained on the C57BL/6 genetic background. Genotyping to detect desired alleles was performed using Transnetyx assays, excluding mice containing both *Aicda*^Cre^ and *Aicda*^KO^ alleles which were genotyped using primers described in Supplementary Table 5. All experimental mice were between 8 and 14-weeks old at the start of experiments. Experiments were designed to use age- and sex-matched mice and, whenever possible, littermates were used as controls. Ctrl mice denote the following genotypes: *Ddx1*^wt/wt^, *Ddx1*^fl/wt^ or *Ddx1*^fl/fl^ and *Cd23*^Cre^ or *Aicda*^Cre^ positive.

### Animal procedures

Mice were immunised by intraperitoneal (I.P) injection with 100μg NP-KLH (Biosearch Technologies) in a 40% (v/v) suspension of Alu-Gel-S (Alum, Universal Biologics) in PBS. Influenza A infection was induced by intranasal administration of A/Hong Kong/1/1968/x31 (x31, H3N2) strain in 30μL PBS at 1×10^4^ colony forming units (CFU), under inhaled isoflurane anaesthesia. To characterise cell-cycle kinetics, immunised mice received I.P. injections of 1mg EdU (Thermo Scientific) and 1.5h later of 2mg BrdU (Sigma Aldrich) prepared in 200μL PBS; spleens were harvested 2.5h after EdU injection. To characterise mRNA translation, immunised mice were injected I.P. with 500μg puromycin (Sigma Aldrich) prepared in 200μL PBS; spleens were harvested 1h after injection. Mice administered with 200μL of PBS were used as negative controls for puromycin labelling. Administration of substances, as well as bodyweight measurements to track Influenza A disease progression, were performed by a technician blind to the genotype. Whenever possible, mice with different genotypes were randomised between cages.

### Flow cytometry

Single-cell suspensions were prepared by mechanical disruption of the spleen through 70μm cell strainers in FACS buffer (PBS supplemented with 0.5% Bovine Serum Albumin (BSA) (Sigma Aldrich) and 2mM EDTA). Alternatively, following HKx31 H2N3 Influenza A infections single-cell suspensions from spleen and mesenteric lymph nodes were prepared in RPMI 1640 media (Sigma Aldrich) supplemented with 10% heat-inactivated Foetal Bovine Serum (hiFBS) (Sigma Aldrich) and 100U Penicillin/Streptomycin (Pen/Strep) (Invitrogen). Cell suspensions were centrifuged at 450xg for 5 minutes at 4°C and treated with ACK lysis buffer (Gibco) at RT for 2 minutes to lyse red blood cells.

For surface staining, cells were incubated with purified anti-mouse CD16/CD32 (BD Biosciences) in FACS buffer for 10 minutes at 4°C, followed by relevant antibody mixes for 1 hour at 4°C. Staining for NP-PE (Biosearch Technologies) and BV605 anti-Mouse IgG1 (BD Biosciences) was performed for each marker individually for 30 minutes at 4°C prior to remaining antibodies. For detection of biotinylated reagents, cells were incubated with streptavidin for 30 minutes at 4°C. To enable exclusion of dead cells, cell pellets were washed with PBS and incubated with Fixable Viability Dyes Near-IR (Thermo Fisher Scientific) or efluor780 (eBiosciences) for 15 minutes at 4°C.

For intracellular staining, cells were fixed and permeabilised using the BD Cytofix/Cytoperm Kit (BD Bioscience) according to manufacturer instructions. Cells were incubated with purified anti-mouse CD16/CD32 in 1x BD Perm/Wash buffer for 10 minutes at 4°C, followed by relevant antibodies overnight at 4°C. For staining that requires secondary antibody detection, cells were washed and stained with relevant antibodies in 1x BD Perm/Wash buffer 1-2 hours at RT.

Samples were acquired using a BD LSR Fortessa (BD Biosciences) or Cytek Aurora Spectral (Cytek Biosciences) analysers. Samples were sorted using a BD FACS ARIA III or Thermo Fisher Scientific BigFoot. Fluorescence compensation or spectral unmixing was performed utilising single-colour stained cells or Anti-Rat and Anti-Hamster Ig κ /Negative Control Compensation Particles Set (BD Biosciences) with BD FACSdiva (v9.0) or SpectralFlo (v3.0) software, respectively. Data was analysed using FlowJo software (TreeStar, v10.8.2).

### Immunofluorescence

Splenic sections were treated as described in Fra-Bido et al., 2021, with minor modifications. The following primary antibodies were used: biotin anti-mouse CD21/35 (clone 8D9), AF488 anti-mouse Ki67 (clone SolA15), AF700 anti-mouse IgD (clone 11-26c.2a) and unconjugated anti-mouse CD3 (clone 500A2); and for secondary detection AF647-conjugated Streptavidin (BioLegend) and AF568-conjugated goat anti-hamster IgG (Life Technologies) were used. After staining, splenic sections were mounted in VectaShield Vibrance Antifade Mounting Medium with DAPI (2Bscientific). Images were acquired using the Leica Stellaris microscope with 20x and 40x objectives and the LAS X office software (Leica Microsystems, v1.4.5); 2-4 non-consecutive sections were assessed per spleen with 3-4 mice per group.

Image processing and quantification was performed utilising an automated pipeline developed by Babraham Imaging Facility (WhiSRC, https://github.com/hannekeo/WhISCR) utilising CellProfiler (v4.2.4)^71^ and ImageJ (FIJI) (v2.1.0)^72^. Full splenic sections were cut into tiles (630 by 630 pixels) for image processing and individual cells were identified in each tile based on DAPI nuclear stain using STARDist program in Image J^73^. The LZ area was defined on groups of CD21/CD35^+^ cells (minimum 20 individual cells) identifying the FDC network; the DZ area was defined on Ki67^+^CD35^-^ clusters (minimum 15 cells) within the GC B-cell population. DZ and LZ areas were combined to define total GC area (minimum 35 cells). If only a Ki67^+^CD35^-^ DZ area was identified, the putative GC structure was excluded. In GCs mostly comprised of Ki67^-^CD21/35^+^ LZ area only, the GC quantification was excluded if the number of Ki67^+^ cells identified within the total GC structure was smaller than 25. This was to ensure GCs were identified in a model which resulted in loss of DZ GC B-cells.

### NP-specific ELISA

ELISA plates (Nunc-ImmunoTM, Thermo Fisher Scientific) were coated with capture reagent in PBS overnight at 4°C. Plates were washed with wash buffer (PBS supplemented 0.05% (v/v) Tween) and blocked with 1% (w/v) BSA in PBS for 1 hour at RT to prevent non-specific binding. Plates were washed prior to the addition of serial dilutions of serum samples in PBS supplemented with 0.1% (w/v) BSA and incubated for 2 hours at RT. NP-specific IgM antibodies were detected by addition of NP-BSA-Biotin for 2 hours and streptavidin-HRP for 20 minutes. NP-specific IgG1 antibodies were detected by addition of HRP-anti mouse IgG1 for 1 hour. Plates were developed with TMB solution (Biolegend) for up to 4 minutes, when the reaction was stopped with 0.5 M H_2_SO_4_. The absorption was measured at 450 nm using the PHERAstar FD microplate reader (BMG Labtech) with PHERAstar data analysis software v3.32. End-point titres were extrapolated from serum dilution curves with arbitrary 0.2 optical density value (NP-specific IgM) or 3x background values obtained from non-immunised controls (NP-specific IgG1) utilised as threshold for each individual mouse at varying timepoints post-immunisation.

### J_H_4 intron sequencing

Genomic DNA was isolated from IgG1^+^ GC B-cells (B220^+^CD19^+^CD38^lo^CD95^+^) sorted from spleens of immunised mice. For amplification of the J_H_4 intron, the protocol published in Sible et al., 2021^26^ was used, with minor modifications. In brief, external PCR was performed utilising 100ng of genomic DNA and Phusion HF polymerase (ThermoScientific) as per manufacturer’s instructions, at a final volume of 20μL per reaction. Samples were PCR amplified (98°C 3’; 21 cycles (98°C 30’’, 55°C 30’’, 72°C 90’’); 72°C 5’) and 1μL of reaction product was used for PCR with internal primers (98°C 3’; 21 cycles (98°C 30’’, 55°C 30’’, 72°C 30’’); 72°C 5’). PCR amplicons were ligated into the TOPO-TA vector (Invitrogen) and transformed into NEB-10b chemically competent E. coli (New England Biolabs) according to the manufacturer’s protocol. Colony PCR was performed utilising M13 forward and reverse primers, bands >600bp were submitted for Sanger sequencing. Sequences were aligned using SeqMan Pro (DNASTAR) and analysed for nucleotide substitutions. The number of mutations per sequence alongside the specific nucleotide change and location was quantified.

### VDJ-Igh1 sequencing

Individual MYC-eGFP^+^IgG1^+^ GC B-cells (Lin^-^B220^+^CD19^+^CD38^lo^CD95^+^) were sorted into 96-well plates containing 10μL lysis buffer (10mM Tris-HCL, 1mM EDTA pH7.4 buffer with 2U/μL RNAsin Plus (Promega), 5mM Dithiothreitol (DTT) and 1% NP40). For cDNA synthesis, RNA was incubated with 2.5ng/μL of Random Hexamer primers (Invitrogen) at 65°C for 5 minutes, followed by Superscript IV (Invitrogen) (23°C for 10 minutes, 50°C for 30 minutes, 80°C for 10 minutes) in a final reaction volume of 25μL.

For amplification of the VDJ region, nested PCR was adapted from protocol published in Von Boehmer et al., 2016^33^. In brief, nested PCR was performed utilising 1-2μL of cDNA per well with GoTaq Hot Start Polymerase and 0.3μM of external forward primers mixture in equal proportions and 0.2μM of external reverse primer, at a final volume of 50μL per reaction. Samples were PCR amplified (94°C 5’; 45 cycles (94°C 30’’, 46°C 30’’, 72°C 55’’); 72°C 10’) and 5μL of reaction product was used for PCR with internal primers (94°C 5’; 40 cycles (94°C 30’’, 57°C 30’’, 72°C 50’’); 72°C 10’). PCR products were purified and submitted for Sanger sequencing. Sequences were aligned to the IMGT mouse heavy chain gene database using NCBI IgBlast to identify V, D and J sequences utilised and number of mutations compared to germline sequences. Singletons and expanded clones were inferred from V(D)J sequences and based on identical IGHV and IGHJ gene annotations, and the length of the junction region.

### Single-cell RNA sequencing

Individual MYC-eGFP^+^ GC B-cells (Lin^-^B220^+^CD19^+^CD38^lo^CD95^+^) were sorted using Thermo Fisher Scientific BigFoot into 96-well plates containing 2μL lysis buffer (0.2% Triton with 1U/μL SUPERNasin (Invitrogen)). Samples were subjected to the Smart-seq2 protocol described in Picelli et al., 2014^74^. Amplification of cDNA was performed using 21 cycles of 98°C 20’’, 67°C 15’’, 72°C 6’. A maximum concentration of 0.2ng/μL cDNA was used for library construction using a miniaturised (1/12.5 volumes) Nextera XT (Illumina) protocol with custom 8bp dual index primers at 0.5μM. Resulting libraries were pooled in batches of 384 and cleaned up using AMPure XP Beads before spiking in 1% Illumina phiX Control v3. Libraries were sequenced on NovaSeq 6000 (Illumina) with NVCS v1.7.5 and RTA v3.4.4 software, set up for 150bp pair-end reads. The data was demultiplexed and converted to fastq using bcl2fastq2.

Raw fastq sequence files were trimmed and underwent quality control utilising Trim Galore (https://doi.org/10.5281/zenodo.7598955). Trimmed sequences were then aligned to mouse genome (Mus musculus GRCm39_v103) utilising HISAT v2.2.1.^75^. Genome-mapped BAM files were processed through SeqMonk (v1.48.2) for gene quantification. The matrix of gene counts was then used as input for analysis by the R package Seurat (v3.1.4.)^76^. Additionally, gene counts were complexed with index sorted data for CXCR4, CD86, CD83 and MYC-eGFP. The dataset was normalised by using the centred log ratio transformation and cells containing more than 20% of sequence reads aligned to mitochondrial genes were excluded prior to normalisation. Next, single cells were clustered with the Seurat workflow and their gene expression evaluated. Assignment of single cells in cell cycle phases was performed utilising the CellCycleScoring function. DZ and LZ definition was based on markers defined in Victora et al 2012^37^ and added utilising the AddModuleScore function. To enable Gene Set Enrichment Analysis (GSEA), pseudobulk analysis using global normalisation (RPM) function was performed for each group. GSEA was performed using the Mouse_GOBP_AllPathways from Merico et al., 2010^41^.

### B-cell cultures

Splenic cell suspensions were prepared in RPMI 1640 media supplemented with 10% hiFBS and 100U Pen/Strep as described above. B-cells were enriched to >95% purity by magnetic depletion using a B-cell isolation kit as per manufactures instructions (Miltenyi Biotech). B-cells were cultured at 5×10^6^ cells per mL in Iscove’s Modified Dulbecco’s Medium (IMDM) (Invitrogen) supplemented with 10% hiFBS, 100U/mL Pen/Strep and 50μM b-Mercaptoethanol (LifeTech). B-cell stimulation was achieved with 20μg/mL F(ab’)₂ Fragment Goat anti-mouse IgM μ-chain specific (Jackson ImmunoResearch) in the presence of 20ng/mL recombinant murine IL4 (Peprotech).

For analysis of mRNA translation, stimulated B-cells were pulsed with 12μg/mL of puromycin (Sigma) at designated timepoint for 15 minutes prior to cell harvest. For a negative control, additional wells were treated with 10μM cycloheximide (Sigma) for 5 minutes, prior to puromycin pulse. Puromycin incorporation was detected by intracellular staining with AF488 anti-Puromycin antibody (Merck, MABE343-AF488) overnight at 4°C.

### Polysome fractionation

Stimulated B-cells were pulsed with 100μg/mL of cycloheximide for 5 minutes prior to cell harvest and lysed on ice for 15 minutes in lysis buffer (15mM Tris-HCL pH 7.5, 300mM NaCL, 15mM MgCl_2_, 100μg/mL cycloheximide and 1000U RNaseOUT (Invitrogen)) plus 1% Triton X-100. Lysates were cleared by centrifugation at 10000g for 20 minutes and layered on to 10-50% sucrose gradients prepared in lysis buffer. Gradients were subjected to centrifugation at 32,000 rpm in a Sorvall SW32.1Ti rotor for 3 hours at 4°C. Fractions were collected from the top of the gradient using a BioComp gradient fractionation system with continuous monitoring of the absorbance at 254 nm.

### Northen blot analysis

Total RNA was isolated from stimulated B-cells and resolved on a denaturing 8% PAGE gel containing 6M urea in 1xTBE at 300V. RNA was transferred to a Hybond N+ membrane in 0.5xTBE at 30V for 1 hour and 30 minutes. Following UV crosslinking (120 J/cm^2^), membranes were blocked in Church hybridisation buffer at 42°C for 30 minutes. This was followed by incubation with 100ng of 5’-end biotin-labelled tyrosine-tRNA probe and IRDye 680RD-labelled proline-tRNA probe overnight at 42°C. Unspecific binding was removed by washing twice with 1xSSC buffer, including 1% SDS for increased stringency. For detection of biotinylated probes, membranes were incubated with IRDye 800RD-labelled streptavidin for 15 minutes at room-temperature. Membranes were visualised in a LiCor Odyssey Gel Scanner and relative quantification of tRNA bands was done using ImageJ (FIJI) (v2.10).

## Data availability

The scRNA-Seq data on MYC-eGFP^+^ GC B-cells we generated in this study is available under the GEO accession number GSE278507. The bulk RNA-seq data on GC B-cell subsets used here is deposited under the GEO accession number GSE278742.

## Code availability

The code used for processing and quantification of confocal images is available at: https://github.com/hannekeo/WhISCR.

## Notes

### Competing Interest Statement

The authors have declared no competing interest.

## References

1. Victora, G.D. & Nussenzweig, M.C. Germinal Centers. Annu Rev Immunol 40, 413–442 (2022).

2. Victora, G.D. et al. Germinal center dynamics revealed by multiphoton microscopy with a photoactivatable fluorescent reporter. Cell 143, 592–605 (2010).

3. Bannard, O. et al. Germinal center centroblasts transition to a centrocyte phenotype according to a timed program and depend on the dark zone for effective selection. Immunity 39, 912–924 (2013).

4. Calado, D.P. et al. The cell-cycle regulator c-Myc is essential for the formation and maintenance of germinal centers. Nat Immunol 13, 1092–1100 (2012).

5. Dominguez-Sola, D. et al. The proto-oncogene MYC is required for selection in the germinal center and cyclic reentry. Nat Immunol 13, 1083–1091 (2012).

6. Nakagawa, R. et al. Permissive selection followed by affinity-based proliferation of GC light zone B cells dictates cell fate and ensures clonal breadth. Proc Natl Acad Sci U S A 118 (2021).

7. Gitlin, A.D. et al. HUMORAL IMMUNITY. T cell help controls the speed of the cell cycle in germinal center B cells. Science 349, 643–646 (2015).

8. Gitlin, A.D., Shulman, Z. & Nussenzweig, M.C. Clonal selection in the germinal centre by regulated proliferation and hypermutation. Nature 509, 637–640 (2014).

9. Finkin, S., Hartweger, H., Oliveira, T.Y., Kara, E.E. & Nussenzweig, M.C. Protein Amounts of the MYC Transcription Factor Determine Germinal Center B Cell Division Capacity. Immunity 51, 324–336.e325 (2019).

10. Long, Z., Phillips, B., Radtke, D., Meyer-Hermann, M. & Bannard, O. Competition for refueling rather than cyclic reentry initiation evident in germinal centers. Sci Immunol 7, eabm0775 (2022).

11. Ersching, J. et al. Germinal Center Selection and Affinity Maturation Require Dynamic Regulation of mTORC1 Kinase. Immunity 46, 1045–1058.e1046 (2017).

12. Pae, J. et al. Cyclin D3 drives inertial cell cycling in dark zone germinal center B cells. J Exp Med 218 (2021).

13. Chou, C. et al. The Transcription Factor AP4 Mediates Resolution of Chronic Viral Infection through Amplification of Germinal Center B Cell Responses. Immunity 45, 570–582 (2016).

14. Inoue, T. et al. The transcription factor Foxo1 controls germinal center B cell proliferation in response to T cell help. J Exp Med 214, 1181–1198 (2017).

15. Sander, S. et al. PI3 Kinase and FOXO1 Transcription Factor Activity Differentially Control B Cells in the Germinal Center Light and Dark Zones. Immunity 43, 1075–1086 (2015).

16. Dominguez-Sola, D. et al. The FOXO1 Transcription Factor Instructs the Germinal Center Dark Zone Program. Immunity 43, 1064–1074 (2015).

17. Kennedy, D.E. et al. Novel specialized cell state and spatial compartments within the germinal center. Nat Immunol 21, 660–670 (2020).

18. Screen, M. et al. RNA helicase EIF4A1-mediated translation is essential for the GC response. Life Sci Alliance 7 (2024).

19. Ribeiro de Almeida, C., et al. RNA Helicase DDX1 Converts RNA G-Quadruplex Structures into R-Loops to Promote IgH Class Switch Recombination. Mol Cell 70, 650–662.e658 (2018).

20. Popow, J., Jurkin, J., Schleiffer, A. & Martinez, J. Analysis of orthologous groups reveals archease and DDX1 as tRNA splicing factors. Nature 511, 104–107 (2014).

21. Jurkin, J. et al. The mammalian tRNA ligase complex mediates splicing of XBP1 mRNA and controls antibody secretion in plasma cells. EMBO J 33, 2922–2936 (2014).

22. Lam, J.H. & Baumgarth, N. The Multifaceted B Cell Response to Influenza Virus. J Immunol 202, 351–359 (2019).

23. Cumano, A. & Rajewsky, K. Structure of primary anti-(4-hydroxy-3-nitrophenyl)acetyl (NP) antibodies in normal and idiotypically suppressed C57BL/6 mice. Eur J Immunol 15, 512–520 (1985).

24. Zuccarino-Catania, G.V. et al. CD80 and PD-L2 define functionally distinct memory B cell subsets that are independent of antibody isotype. Nat Immunol 15, 631–637 (2014).

25. Tomayko, M.M., Steinel, N.C., Anderson, S.M. & Shlomchik, M.J. Cutting edge: Hierarchy of maturity of murine memory B cell subsets. J Immunol 185, 7146–7150 (2010).

26. Sible, E., Zheng, S., Choi, J.E. & Vuong, B.Q. Analysis of Somatic Hypermutation in the JH4 intron of Germinal Center B cells from Mouse Peyer’s Patches. J Vis Exp (2021).

27. Zaheen, A. et al. AID constrains germinal center size by rendering B cells susceptible to apoptosis. Blood 114, 547–554 (2009).

28. Boulianne, B. et al. AID and caspase 8 shape the germinal center response through apoptosis. J Immunol 191, 5840–5847 (2013).

29. Yewdell, W.T. et al. A Hyper-IgM Syndrome Mutation in Activation-Induced Cytidine Deaminase Disrupts G-Quadruplex Binding and Genome-wide Chromatin Localization. Immunity 53, 952–970.e911 (2020).

30. Huang, C.Y., Bredemeyer, A.L., Walker, L.M., Bassing, C.H. & Sleckman, B.P. Dynamic regulation of c-Myc proto-oncogene expression during lymphocyte development revealed by a GFP-c-Myc knock-in mouse. Eur J Immunol 38, 342–349 (2008).

31. Mayer, C.T. et al. The microanatomic segregation of selection by apoptosis in the germinal center. Science 358 (2017).

32. Stewart, I., Radtke, D., Phillips, B., McGowan, S.J. & Bannard, O. Germinal Center B Cells Replace Their Antigen Receptors in Dark Zones and Fail Light Zone Entry when Immunoglobulin Gene Mutations are Damaging. Immunity 49, 477–489.e477 (2018).

33. von Boehmer, L. et al. Sequencing and cloning of antigen-specific antibodies from mouse memory B cells. Nat Protoc 11, 1908–1923 (2016).

34. Jacob, J. & Kelsoe, G. In situ studies of the primary immune response to (4-hydroxy-3-nitrophenyl)acetyl. II. A common clonal origin for periarteriolar lymphoid sheath-associated foci and germinal centers. J Exp Med 176, 679–687 (1992).

35. Xue, H. et al. Artificial immunoglobulin light chain with potential to associate with a wide variety of immunoglobulin heavy chains. Biochem Biophys Res Commun 515, 481–486 (2019).

36. Allen, D., Simon, T., Sablitzky, F., Rajewsky, K. & Cumano, A. Antibody engineering for the analysis of affinity maturation of an anti-hapten response. EMBO J 7, 1995–2001 (1988).

37. Victora, G.D. et al. Identification of human germinal center light and dark zone cells and their relationship to human B-cell lymphomas. Blood 120, 2240–2248 (2012).

38. Toboso-Navasa, A. et al. Restriction of memory B cell differentiation at the germinal center B cell positive selection stage. J Exp Med 217 (2020).

39. van der Maaten, L. & Hinton, G. Visualizing Data using t-SNE. Journal of Machine Learning Research; 2008. pp. 2579–2605.

40. Love, M.I., Huber, W. & Anders, S. Moderated estimation of fold change and dispersion for RNA-seq data with DESeq2. Genome Biol 15, 550 (2014).

41. Merico, D., Isserlin, R., Stueker, O., Emili, A. & Bader, G.D. Enrichment map: a network-based method for gene-set enrichment visualization and interpretation. PLoS One 5, e13984 (2010).

42. Nagayoshi, Y. et al. Loss of Ftsj1 perturbs codon-specific translation efficiency in the brain and is associated with X-linked intellectual disability. Sci Adv 7 (2021).

43. Seedhom, M.O. et al. Paradoxical imbalance between activated lymphocyte protein synthesis capacity and rapid division rate. Elife 12 (2024).

44. Kroupova, A. et al. Molecular architecture of the human tRNA ligase complex. Elife 10 (2021).

45. Roco, J.A. et al. Class-Switch Recombination Occurs Infrequently in Germinal Centers. Immunity 51, 337–350.e337 (2019).

46. King, H.W., et al. Single-cell analysis of human B cell maturation predicts how antibody class switching shapes selection dynamics. Sci Immunol 6 (2021).

47. Pape, K.A. et al. Visualization of the genesis and fate of isotype-switched B cells during a primary immune response. J Exp Med 197, 1677–1687 (2003).

48. Toellner, K.M., Gulbranson-Judge, A., Taylor, D.R., Sze, D.M. & MacLennan, I.C. Immunoglobulin switch transcript production in vivo related to the site and time of antigen-specific B cell activation. J Exp Med 183, 2303–2312 (1996).

49. Sundling, C. et al. Positive selection of IgG. Immunity 54, 988–1001.e1005 (2021).

50. van Riggelen, J., Yetil, A. & Felsher, D.W. MYC as a regulator of ribosome biogenesis and protein synthesis. Nat Rev Cancer 10, 301–309 (2010).

51. Tan, T.C.J. et al. Suboptimal T-cell receptor signaling compromises protein translation, ribosome biogenesis, and proliferation of mouse CD8 T cells. Proc Natl Acad Sci U S A 114, E6117–E6126 (2017).

52. Marchingo, J.M., Sinclair, L.V., Howden, A.J. & Cantrell, D.A. Quantitative analysis of how Myc controls T cell proteomes and metabolic pathways during T cell activation. Elife 9 (2020).

53. Shaffer, A.L. et al. XBP1, downstream of Blimp-1, expands the secretory apparatus and other organelles, and increases protein synthesis in plasma cell differentiation. Immunity 21, 81–93 (2004).

54. Hu, C.C., Dougan, S.K., McGehee, A.M., Love, J.C. & Ploegh, H.L. XBP-1 regulates signal transduction, transcription factors and bone marrow colonization in B cells. EMBO J 28, 1624–1636 (2009).

55. Taubenheim, N. et al. High rate of antibody secretion is not integral to plasma cell differentiation as revealed by XBP-1 deficiency. J Immunol 189, 3328–3338 (2012).

56. Todd, D.J. et al. XBP1 governs late events in plasma cell differentiation and is not required for antigen-specific memory B cell development. J Exp Med 206, 2151–2159 (2009).

57. Goodarzi, H. et al. Modulated Expression of Specific tRNAs Drives Gene Expression and Cancer Progression. Cell 165, 1416–1427 (2016).

58. Gingold, H. et al. A dual program for translation regulation in cellular proliferation and differentiation. Cell 158, 1281–1292 (2014).

59. Thandapani, P. et al. Valine tRNA levels and availability regulate complex I assembly in leukaemia. Nature 601, 428–433 (2022).

60. Rak, R. et al. Dynamic changes in tRNA modifications and abundance during T cell activation. Proc Natl Acad Sci U S A 118 (2021).

61. Guillon, J. et al. tRNA biogenesis and specific aminoacyl-tRNA synthetases regulate senescence stability under the control of mTOR. PLoS Genet 17, e1009953 (2021).

62. Huh, D. et al. A stress-induced tyrosine-tRNA depletion response mediates codon-based translational repression and growth suppression. EMBO J 40, e106696 (2021).

63. Gao, L. et al. Selective gene expression maintains human tRNA anticodon pools during differentiation. Nat Cell Biol 26, 100–112 (2024).

64. Guimaraes, J.C. et al. A rare codon-based translational program of cell proliferation. Genome Biol 21, 44 (2020).

65. Giguère, S. et al. Antibody production relies on the tRNA inosine wobble modification to meet biased codon demand. Science 383, 205–211 (2024).

66. Liu, Y. et al. tRNA-m. Nat Immunol 23, 1433–1444 (2022).

67. Johnson, P.F. & Abelson, J. The yeast tRNATyr gene intron is essential for correct modification of its tRNA product. Nature 302, 681–687 (1983).

68. van Tol, H. & Beier, H. All human tRNATyr genes contain introns as a prerequisite for pseudouridine biosynthesis in the anticodon. Nucleic Acids Res 16, 1951–1966 (1988).

69. Strobel, M.C. & Abelson, J. Effect of intron mutations on processing and function of Saccharomyces cerevisiae SUP53 tRNA in vitro and in vivo. Mol Cell Biol 6, 2663–2673 (1986).

70. Kwon, K. et al. Instructive role of the transcription factor E2A in early B lymphopoiesis and germinal center B cell development. Immunity 28, 751–762 (2008).

71. Stirling, D.R. et al. CellProfiler 4: improvements in speed, utility and usability. BMC Bioinformatics 22, 433 (2021).

72. Schindelin, J., et al. Fiji: an open-source platform for biological-image analysis. Nat Methods 9, 676-682 (2012).

73. Schmidt, U. et al. Cell Detection with Star-Convex Polygons. Medical Image Computing and Computer Assisted Intervention - Miccai 2018, Pt Ii 11071, 265–273 (2018).

74. Picelli, S. et al. Full-length RNA-seq from single cells using Smart-seq2. Nat Protoc 9, 171–181 (2014).

75. Kim, D., Paggi, J.M., Park, C., Bennett, C. & Salzberg, S.L. Graph-based genome alignment and genotyping with HISAT2 and HISAT-genotype. Nat Biotechnol 37, 907–915 (2019).

76. Stuart, T. et al. Comprehensive Integration of Single-Cell Data. Cell 177, 1888–1902.e1821 (2019).

